# Benchmarking of duplex sequencing approaches to reveal somatic mutation landscapes

**DOI:** 10.64898/2025.12.12.692823

**Authors:** Yang Zhang, Vinayak V. Viswanadham, Michail Andreopoulos, Dominik Glodzik, Ruolin Liu, Lovelace J. Luquette, Se-Young Jo, Azeet Narayan, Muchun Niu, Lisa Anderson, Joseph A. Brew, Hsu Chao, Carrie Cibulskis, Guanlan Dong, Uday S. Evani, William C. Feng, Marta Grońska-Pęski, Adrienne Helland, Nazia Hilal, Nisrine T. Jabara, Hu Jin, Ning Li, Monica D. Manam, Shayna L. Mallett, Alexi Runnels, Constantijn Scharlee, Carter Smith, SMaHT Duplex Sequencing Focus Group, Diane Shao, Christopher A. Walsh, Viktor A. Adalsteinsson, Eunjung Alice Lee, Peter J. Park, Kristin G. Ardlie, Soren Germer, Richard A. Gibbs, Sangita Choudhury, Harsha V. Doddapaneni, Gilad D. Evrony, Chenghang Zong, Tim H.H. Coorens

## Abstract

Detecting somatic mutations in normal tissues is challenging due to sequencing errors and the low allele fractions of post-zygotic variants. Duplex sequencing greatly reduces errors and can detect mutations at any allele fraction, but systematic, cross-platform comparisons are lacking. We present a comprehensive benchmarking of six duplex sequencing technologies used by the SMaHT Network: CODEC, CompDuplex-seq, HiDEF-seq, NanoSeq, ppmSeq, and VISTA-seq. We evaluated their performance using cord blood DNA, a tumor-normal cell line mixture, and homogenates from six human tissues. Each method shows distinct profiles in genomic footprint, sensitivity, and cost. Despite differences in library construction and sequencing platforms, estimates of mutation rates and mutational signatures are highly concordant. Integration with ultra-deep whole-genome sequencing shows that duplex approaches sensitively capture mutations and signatures beyond embryonic or clonally expanded variants. These results provide a foundation for selecting duplex methods and interpreting their data, enabling scalable single-molecule analyses of somatic mutation landscapes.

**Graphical Abstract:** 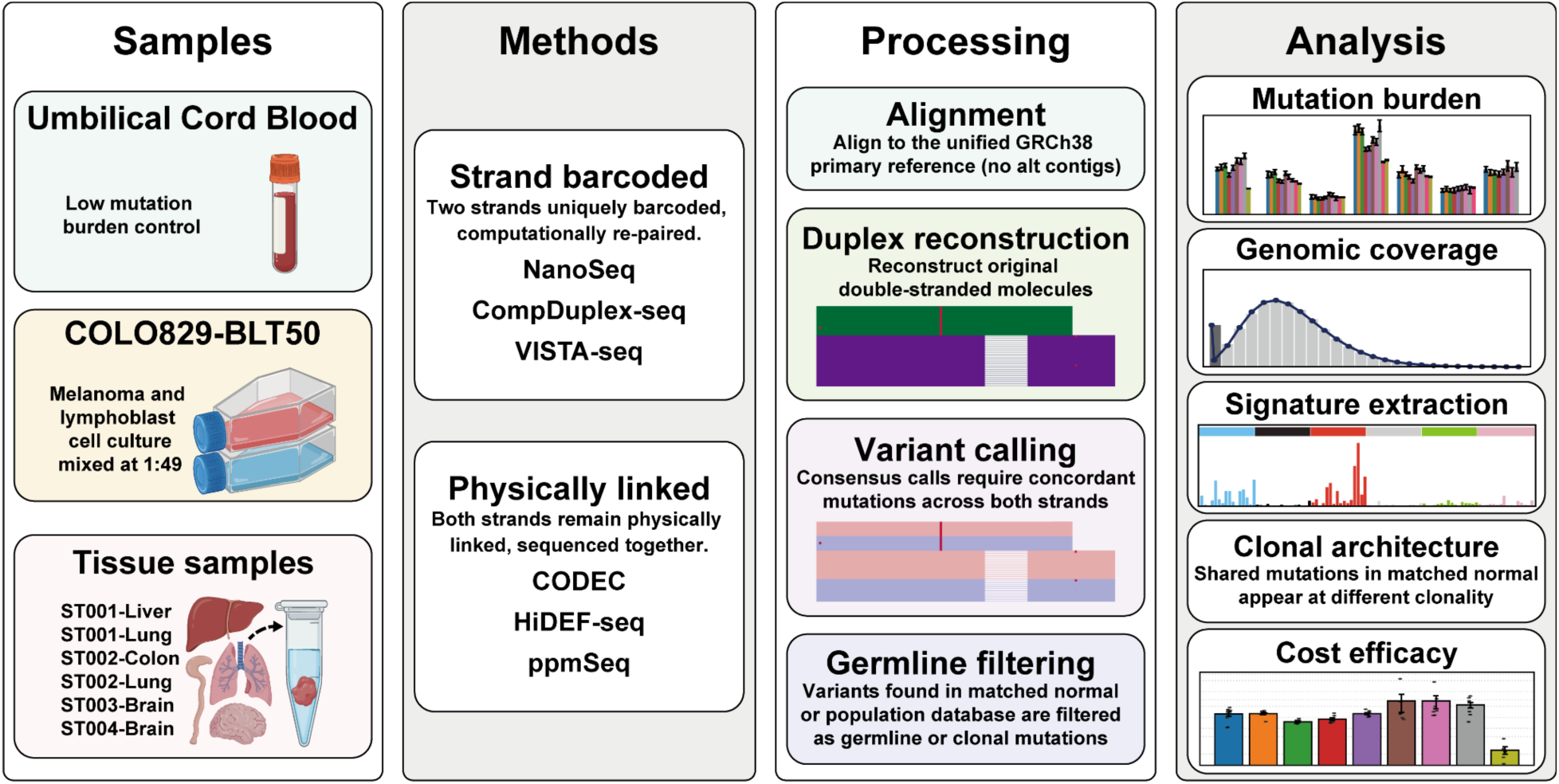

## Main

Somatic mutations accumulate in all cells after fertilization due to both endogenous processes, such as DNA replication errors and oxidative damage, and exogenous exposures. The prevalence and diversity of somatic mosaicism vary widely across human tissues^1^, shaped by differences in cell type composition and physiology, rates of cell division, and exposure to mutagens. Pathogenic somatic mutations have been implicated in cancer^2^, neurodegenerative disorders^3^, and developmental disorders^4,5^, with their effects modulated by the timing, cell types, and locations in the body in which they occur. Therefore, accurate and efficient profiling of somatic mutations is crucial to understanding their impact in both health and disease. To advance this understanding, the Somatic Mosaicism across Human Tissues (SMaHT) Network^6^ has been developing and applying diverse approaches for profiling somatic mutations across tissue types with the goal of providing a comprehensive framework for studying mosaicism at unprecedented resolution.

The detection of somatic mutations is intrinsically challenging: many mutant alleles are rare and present at variant allele fractions (VAFs) well below 1%. At most one DNA sequence read would support such mutations in deep WGS studies. While panel or targeted sequencing can increase resolution by achieving deeper sequencing^7^, it requires foreknowledge of mutated loci or selection of candidate genes, precluding unbiased, genome-wide analyses. With conventional short-read sequencing, per-base error rates on the order of 10⁻³ overwhelm these low-frequency signals, so deeper sequencing does not reliably separate true mutations from artifacts. Direct single-cell sequencing^8–10^ or sequencing of single-cell clonal expansions^11–14^ can amplify alleles to detectable frequencies but at substantial labor and cost. Duplex sequencing^15,16^ addresses these constraints by dramatically lowering per-base error rates to below the signal level of somatic mutations, such that somatic mutations can be reliably called even when detected in only one molecule. Duplex sequencing reduces the error rate by reading both strands of the same DNA duplex and requiring concordance between complementary strands. Because most artifacts introduced by DNA damage, library preparation, or sequencing do not affect the same locus on both strands simultaneously, duplex consensus suppresses these single-stranded errors. Notably, consensus requires obtaining multiple sequencing reads per DNA molecule, which incurs higher cost per final interrogated base pair than conventional sequencing. A pivotal advance was Nanorate Sequencing (NanoSeq)^17^, which utilizes molecular bar codes to identify strand pairs and non-A dideoxy nucleotides (ddBTPs) to eliminate residual duplex sequencing artifacts that arise from DNA polymerase nick extension during A-tailing. However, duplex sequencing methods have been challenged by sample input requirements, the need for specialized analyses that have yet to converge on best practices, and higher costs relative to conventional sequencing.

Motivated by the need for greater diversity in sample inputs, broader genome coverage, and higher duplex efficiency, new duplex sequencing technologies have recently emerged (**Figure 1**). These methods, CODEC^18^, CompDuplex-seq^19^, HiDEF-seq^20^, ppmSeq^21^, and VISTA-seq (based on META-CS^22^), together with NanoSeq^17^, span two duplex-reconstruction approaches: (i) strand-barcoded duplexes (CompDuplex-seq, NanoSeq and VISTA-seq), in which the two strands are tagged and sequenced in separate clusters and later joined bioinformatically; and (ii) physically-linked duplexes (CODEC and HiDEF-seq specifically and ppmSeq to some extent), in which the two strands are physically linked during library preparation and sequencing so a single sequencing cluster interrogates both strands.

**Figure 1.**
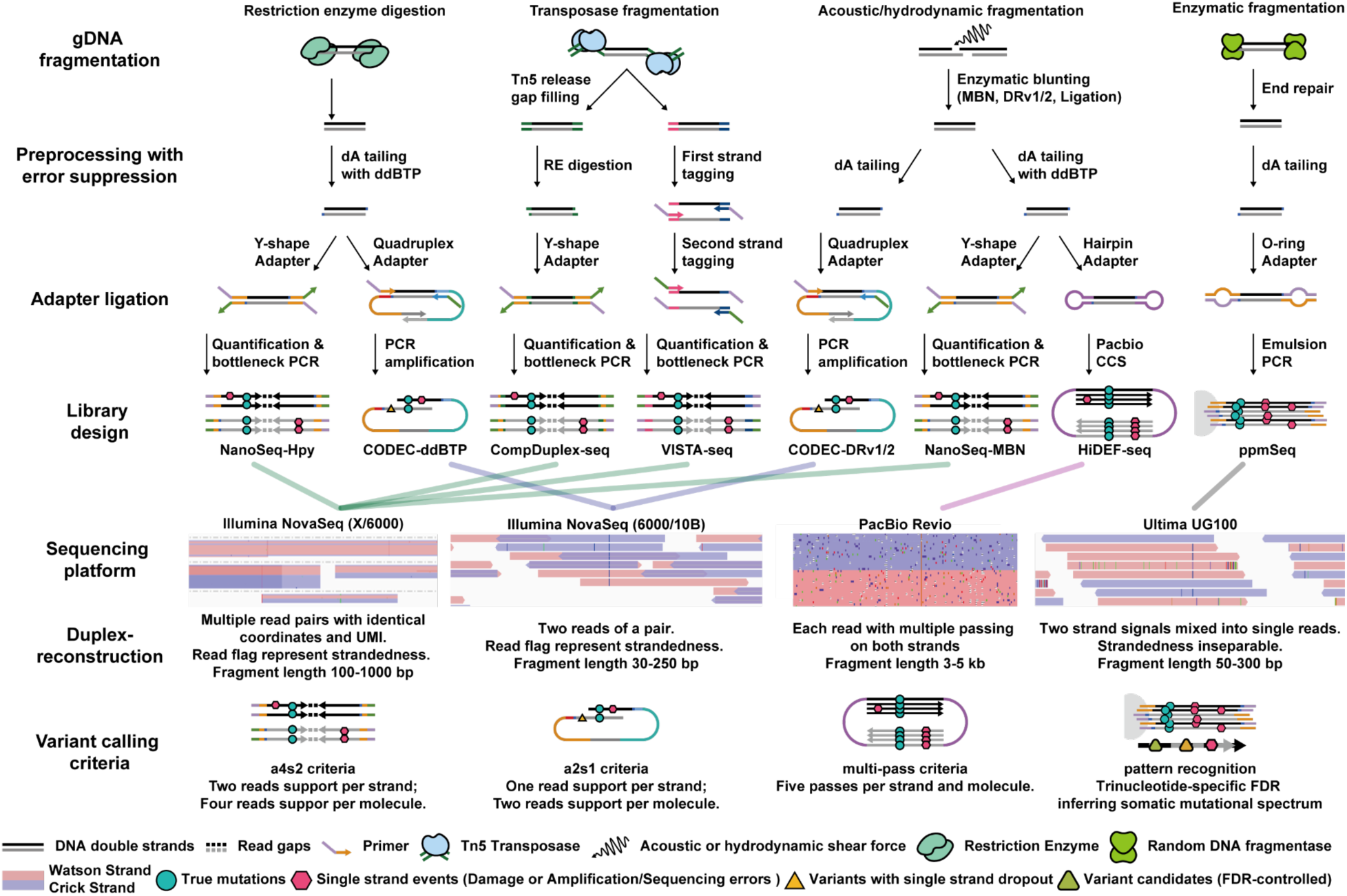
Comparison of library preparation for the six evaluated duplex sequencing methods.

Strand-barcoded duplex sequencing methods, including NanoSeq, CompDuplex-seq, and VISTA-seq, independently barcode the two strands and sequence them as distinct clusters. The complementary strands are then paired *in silico* using alignment coordinates and reverse-complement UMIs. To maximize the probability that both strands are observed, these protocols include a qPCR-calibrated bottlenecking step that limits input molecular complexity to match the intended raw depth. Variants lacking bidirectional support are excluded from consensus, which lowers apparent recovery but strongly suppresses single-strand artefacts and polymerase damage signatures. A major focus of recent work has been expanding genomic reach beyond the original NanoSeq formulation, which was limited to sites created by restriction enzyme cleavage^17,23–25^. Mung Bean NanoSeq (NanoSeq-MBN) optimizes conditions for DNA fragmentation using sonication, which provides random genome-wide coverage, and mung bean nuclease blunting of the resulting single-strand overhangs. VISTA-seq builds on META-CS single-cell duplex sequencing^22^ that uses random combinations of 16 distinct Tn5 transposase sequences to encode strand identity, after which adapters are appended through sequential primer annealing and extension reactions. VISTA-seq enables mutational profiling not only from single-cell input, as in the original META-CS, but also from bulk and micro-bulk input DNA (≈50 nuclei). In CompDuplex-seq, genomic DNA is fragmented in a largely random fashion with minimal Tn5 insertion bias, and the transposase recognition sites are subsequently cut by a restriction enzyme to create sticky ends for adapter ligation, therefore, there is no need for using multiple Tn5 transposase sequences for DNA tagmentation as in the META approach^22^. Because Tn5 tagmentation is mechanically mild relative to acoustic shearing, it helps minimize shear-/oxidation-related DNA damage while maintaining complexity.

Physically-linked duplex sequencing methods, including CODEC, HiDEF-seq, and ppmSeq, instead keep the two strands of each original duplex physically associated during library construction and sequencing, with the goal of boosting duplex reconstruction efficiency. CODEC uses a quadruplex adapter system to concatenate both strands into a single sequencing unit so that the two strands of the input DNA molecule manifest as the two reads of a read-pair. HiDEF-seq employs PacBio single-molecule sequencing with hairpin adapters that concatenates both strands into circularized molecules that can be sequenced multiple times by rolling-circle sequencing. By selecting shorter inserts (≈3–5 kb) and applying nick ligation, HiDEF-seq increases the number of independent sequences per strand and eliminates residual single-strand artifacts to achieve high consensus fidelity; we also introduce here a new version of HiDEF-seq with whole-genome coverage. ppmSeq co-encapsulates both strands within the same emulsion bead, enabling simultaneous amplification and co-sequencing; single clusters thus contain a mixture of signals from both strands. Inconsistent bases (e.g., from damage or amplification errors) manifest as lower base-quality scores, whereas concordant positions score higher. Because practical quality thresholds can drift, per-site genotype calls in ppmSeq become challenging at ultra-low VAF. To address this, ppmSeq analyzes trinucleotide-context-specific false discovery rates (FDRs) to recover accurate mutational spectra.

Together, these technologies span the full spectrum of duplex sequencing approaches, including diverse library preparation methods and sequencing platforms (Illumina, PacBio, and Ultima platforms). Yet a side-by-side, equal-footing assessment of their strengths and limitations is lacking. Therefore, within the SMaHT Network, we established a harmonized, multi-institution study of these six duplex sequencing methods using unified biological materials and standardized performance criteria. This work delivers a practical, application-oriented reference: a common dataset, rigorously comparable metrics, and clear operating points that enable investigators to choose the right assay for discovery screens, mutational-spectrum profiling, or high-confidence site-level calling, while illuminating the trade-offs among breadth, fidelity, recovery efficiency, and cost. It also outlines a generalizable approach for evaluating future duplex sequencing methods.

## Results

### Benchmarking sample design

We designed a harmonized benchmark workflow (**Graphical abstract**) spanning six duplex-sequencing technologies (NanoSeq, CompDuplex-seq, VISTA-seq, CODEC, HiDEF-seq, and ppmSeq) to compare performance on equal footing while preserving each method’s strengths. We included in the study two versions of NanoSeq, one utilizing HpyCH4V restriction enzyme fragmentation (NanoSeq-Hpy) and another utilizing random fragmentation followed by mung bean nuclease end repair (NanoSeq-MBN). For CODEC, we included three versions utilizing different end repair methods (ddBTP^18^, DRv1^26^, and DRv2^27^). To minimize protocol bias, we standardized biological inputs and sample handling across sites and coordinated execution with developer groups so that each team constructed libraries and conducted method-specific primary processing through variant calling under fully optimized conditions. All downstream evaluations used a centralized, method-agnostic pipeline. We report general, application-relevant metrics across the workflow in **Table S1**, with key performance metrics presented as a graphical table in **Figure 2A**.

**Figure 2.**
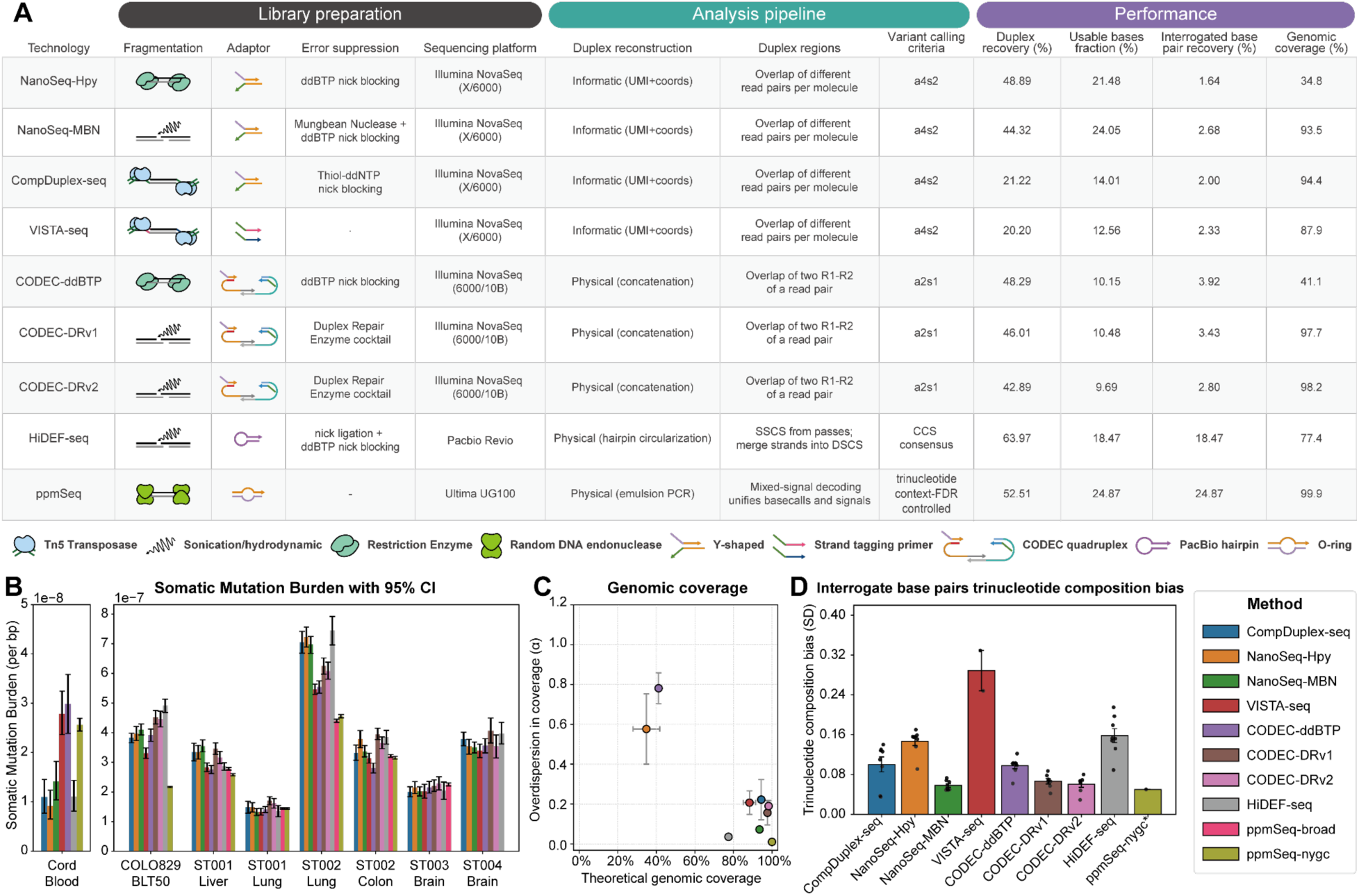
General metrics A. Summary table for mean performance of benchmarked methods across evaluated samples. The legend below the table shows fragmentation and adapter approaches. B. Estimated somatic mutation burden, calculated as number of mutations per final interrogated base pair. C. Scatter plot showing genome coverage and evenness based on a zero-inflated negative binomial model of read depth across the genome (Methods). D. Library-specific trinucleotide composition bias at interrogated base pairs (SD).

To ensure robust, application-oriented contrasts, we profiled three distinct sample types, each of which assesses specific duplex sequencing performance metrics. A cord-blood control provided a low-burden reference to benchmark specificity and background error rates. A COLO829 cell-line mix (1:49 tumor:normal) with a matched normal supplied a set of known variants for estimating sensitivity, limit of detection, and precision–recall behavior. Finally, homogenates from six human tissues (**Table S1**) modeled real-world heterogeneity to stress-test coverage evenness, duplex recovery, and spectrum stability.

### Somatic mutation burden estimation

Our primary readout is each method’s ability to recover the somatic mutation burden, defined as the number of somatic mutations observed divided by the effective depth (interrogated base pairs (bp), after all filters are applied), which can also be expressed as the number of mutations per diploid cell equivalent. In the low-burden cord-blood sample (**Figure 2B**), NanoSeq-Hpy, NanoSeq-MBN, CompDuplex-seq, and HiDEF-seq measured ∼1×10⁻⁸ mutations per bp, specifically: 56.9 (95% CI 40.2–76.3), 87.4 (64.2–112.4), 67.8 (47.4–89.8), 67.9 (50.2-88.5) mutations per cell, respectively, which is concordant with prior studies^17,20^. In contrast, VISTA-seq, CODEC-ddBTP, and ppmSeq measured ∼2.6×10⁻⁸ mutations per bp, specifically: 172.1 (147.0–200.4), 184.4 (147.5–221.3), and 158.1 (150.4–166.1) mutations per cell, respectively. Because cord blood harbors very few true mutations, single-digit count fluctuations can increase variability in burden estimates, potentially alongside subtle differences in sample quality across centers and bioinformatic pipelines. Nevertheless, burdens largely remain within the 10⁻⁸ scale. Meanwhile, the consistent estimation by four methods (NanoSeq-Hpy, NanoSeq-MBN, CompDuplex-seq, and HiDEF-seq) and concordance with other methods^12,28^ reflect a convergence to the ground truth. It is worth noting that when a higher-burden context (≈10⁻⁷-10⁻⁶) is used for comparison as detailed below, all the methods converge to consistent estimates across samples.

In the COLO829 tumor mix (**Figure 2B**), most methods measured a substitution burden of ∼3.5–4.0×10⁻⁷. An exception is ppmSeq, which reports roughly half that value. This downward shift reflects its calling paradigm: instead of using a standalone matched normal to filter germline and clonal variants (as done uniformly by the other methods with the COLO829 BL line), ppmSeq infers which mutations are somatic, occasionally leading to misassignments. The ppmSeq mutation calling workflow uses trinucleotide–context false discovery rate (FDR) from germline and clonal signals within the same data and then applies pattern recognition to classify somatic candidates. As a result, many bona fide tumor variants are thus re-labeled as clonal and removed, systematically underestimating the somatic burden in this sample type. This result indicates that overall accuracy level of ppmSeq is on the scale of 10⁻⁷ per base. Further development for variant calling of ppmSeq is actively underway. Across the remaining tissues, methods are largely concordant, yielding consistent burden estimations across tissues despite differences in chemistry and read structure. One notable outlier is ST002–Lung, where between-method dispersion is larger, for example, when compared with ST001-Lung. The two brain donors (ST003 and ST004) are age-matched to the other tissue donors (ST003, 22 years; 1309.2 somatic mutations per cell) < (ST004, 73 years; 2258.5 per cell), averaging 18.6 mutations per cell per year across methods. Together, these patterns indicate that exposure history, tissue type, and age rather than platform dominate variability. Overall, the six duplex platforms yield concordant estimates while delineating clear tissue-specific differences in somatic mutation burden.

### Genomic coverage breadth and uniformity

We next compared genome breadth and coverage evenness. Duplex-sequencing methods differ markedly in how much of the genome they interrogate and how uniformly they distribute depth, properties affected by DNA fragmentation (restriction enzymes versus mechanical fragmentation versus transposase fragmentation) and downstream library construction.

Under ideal random sampling, per-base coverage would be Poisson-distributed across the genome. However, real samples deviate from this, for example due to biases in fragmentation that create over-dispersion and an excess of regions with zero coverage (e.g., regions lacking restriction sites). To quantify breadth and uniformity in genome coverage without confounding by overall depth, we modeled per-base duplex coverage with a zero-inflated negative binomial (ZINB) model, which simultaneously captures zero inflation and variance inflation. This provided a consistently superior fit to genome coverage relative to a Poisson model across all technologies (**Figure S1**) and enabled depth-normalized comparisons of genome coverage breadth and uniformity.

Overall, genome coverage metrics followed fragmentation strategy. Methods utilizing non-random enzymatic fragmentation (NanoSeq-Hpy and CODEC-ddBTP) exhibited narrower genomic breadth (30-40%) and elevated variance (**Figure 2C**), consistent with sequence-specific cutting and post-fragmentation inefficiencies. In contrast, methods utilizing random fragmentation such as sonication fragmentation, hydrodynamic fragmentation, random enzymatic fragmentation, or Tn5-transposase fragmentation (NanoSeq-MBN, CompDuplex-seq, CODEC-DRv1/DRv2, HiDEF-seq, VISTA-seq, and ppmSeq) approached Poisson expectations with broader, more uniform coverage. Notably, breadth values <100% primarily reflect excluded masked regions rather than incomplete sampling. Coverage uniformity has direct downstream implications for mutational-signature analysis because uneven genomic representation can distort the observed trinucleotide context frequencies.

### Trinucleotide composition bias

Interrogated-base composition was quantified by binning covered bases into 32 trinucleotide classes and comparing each library to hg38 baseline frequencies under the same genome mask. Bias was summarized as the standard deviation (SD) across contexts (**Figure 2D**). Lower SD indicates closer agreement with the genome; higher SD reflects stronger composition skew, independent of direction.

Whole-genome profiling methods (NanoSeq-MBN, CODEC-DRv1/2, and ppmSeq) show the smallest SDs, consistent with their near-uniform genomic coverage reported above; methods with more uneven coverage generally exhibit larger SDs. Restriction-enzyme-based methods (NanoSeq-Hpy and CODEC-ddBTP) display elevated, though not maximal, composition bias. With a single enzyme, depletion is pronounced at the precise recognition motif, for example in NanoSeq-Hpy versus NanoSeq-MBN (at the HpyCH4V motif TG|CA; **Figure S2C(i)**). Using a mix of enzymes (HpyCH4V and AluI) appears to attenuate depletion (**Figure S2C(ii)**), likely because distinct recognition sites distribute cleavage across many motifs, diluting the scarcity of any one context.

CompDuplex exhibits a low degree bias in high-integrity cell-line DNA, whereas increased bias in tissue homogenates (**Figure 2D**). This difference indicates a potential sequence bias exists in nicks induced by mechanical homogenization. It is worth pointing out that these nicks are largely removed or repaired in NanoSeq-MBN, and CODEC-DRv1/2. VISTA-seq shows comparatively high SDs (**Figure 2D**), plausibly reflecting a heavier amplification bias, where context-dependent amplification efficiencies skew trinucleotide representation. HiDEF-seq achieves relatively uniform overall coverage across the genome, yet the base pairs effectively interrogated display some imbalance in trinucleotide-context representation (**Figure S2D(viii)**). This distortion may in part reflect context-dependent variation in error rates intrinsic to the PacBio sequencing platform.

It is worth noting that, although our data exhibit non-negligible trinucleotide-context bias, the downstream mutational signature analyses remain robust to these distortions, as detailed in the following section.

### Mutational landscapes viewed through different duplex methods

Comparison of mutational profiles across sequencing technologies for a given sample revealed high overall similarity, with the notable exception of ppmSeq mutations supported by multiple reads in tissue homogenates. This discrepancy was not observed in COLO829-BLT50, a sample with a known clonal structure (**Figure S4**). When incorporated into the full mutation catalogue, ppmSeq calls introduced assay-specific components that confounded de novo mutational signature discovery (**Figure S5**). We therefore excluded ppmSeq from the primary signature discovery analysis (**Methods**).

Across the remaining samples and sequencing assays, mutational signature extraction identified five robust signatures (**Figure 3A**), with MuSiCal^29^ (non-negative matrix factorization) and hierarchical Dirichlet process (HDP)^1,30^ mixture modeling independently converging on a highly similar set of signatures (**Figure S6**). Subsequent fitting of these signatures across all samples and sequencing assays revealed no assay-specific signatures; instead, clustering patterns were primarily driven by sample identity, indicating that the *de novo* signatures reflect underlying biological differences between samples rather than technical variation between the duplex sequencing assays (**Figure 3B**).

**Figure 3.**
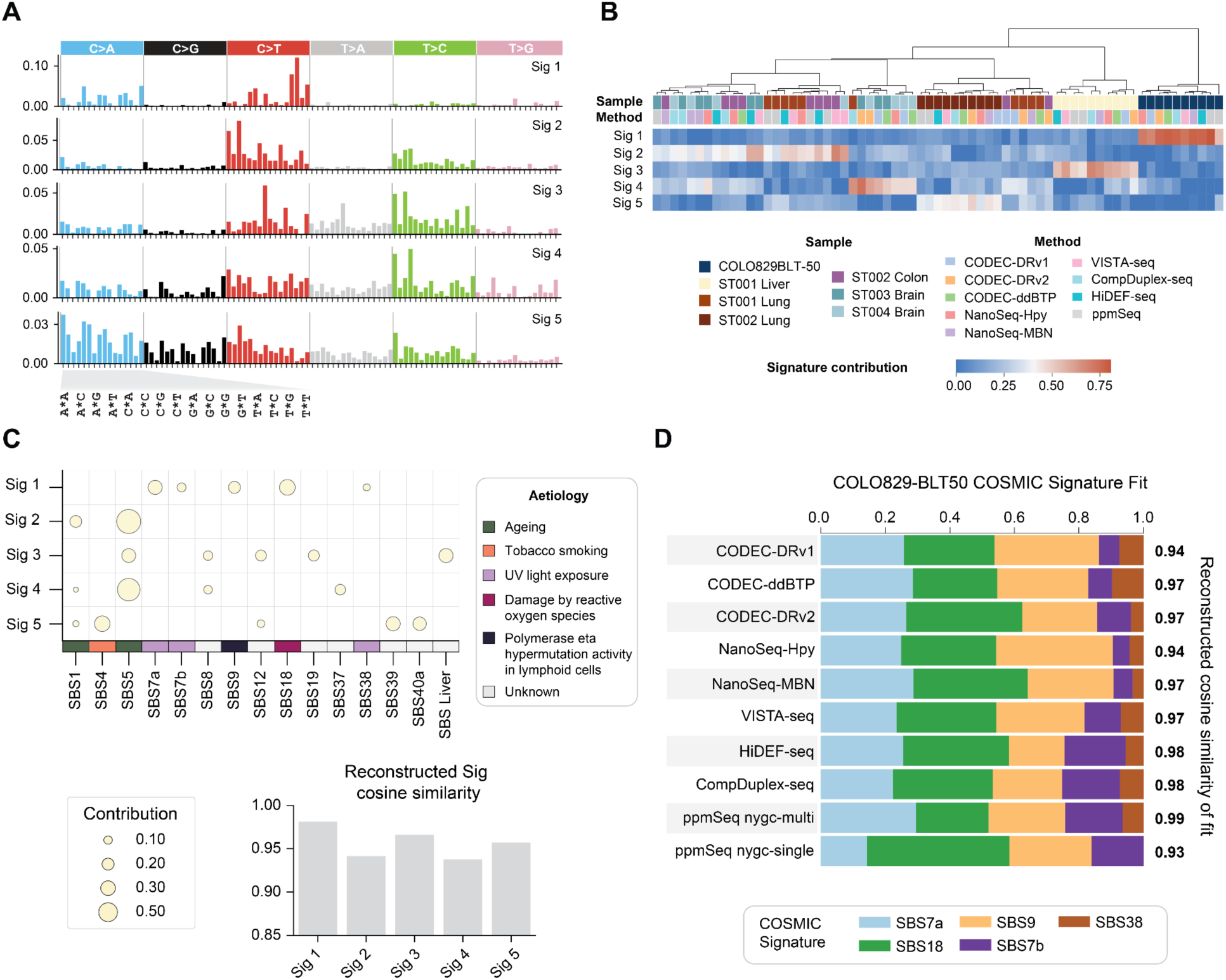
Mutational signature analysis A. De novo mutational signatures extracted using Non-negative Matrix Factorization (NMF) as implemented by MuSiCal. B. Heatmap of signature exposures across samples, hierarchically clustered using Ward’s method to reveal signature exposure patterns by tissue type and sequencing platform. Top rows indicate sample identity, tissue, and sequencing method. C. Decomposition of de novo signatures into known COSMIC signatures, expected for the tissues studied. The size of filled circles indicates the contribution proportion of each COSMIC signature. The accompanying barplot shows cosine similarity between each de novo signature and its reconstruction from the top contributing COSMIC signatures, reflecting how well-known signatures explain the de novo patterns. D. Estimating the abundance of COSMIC signatures SBS7a,7b,9,18,38 detected in de novo “Sig 1” to mutation sets obtained for the COLO829-BLT sample using different duplex sequencing assays. Right: Cosine similarity values of reconstructed profiles.

To further interpret the extracted *de novo* signatures, each was decomposed into known COSMIC reference signatures^31^ (**Figure 3C**). This analysis revealed that tissue-relevant COSMIC single base substitution (SBS) signatures contributed most strongly to the corresponding samples. For instance, in tissue homogenates, the liver sample ST001 exhibited high activity of *de novo* signature 3, which was best explained by the liver-associated COSMIC signature SBS12 and a previously reported liver-specific signature^32^. Similarly, the older lung sample ST002 (74 years old) exhibited an elevated mutation burden (**Figure 2B**) and was enriched for *de novo* signature 5, which was mostly explained by the COSMIC signature SBS4, a signature previously associated with exposure to tobacco smoke^33^.

Analysis of the COLO829-BLT50 mixture sample allowed for a comparison of the detected signatures against the expectation from known cell type composition. The *de novo* signature 1 was decomposed into signatures associated with UV light exposure (SBS7a, SBS7b, SBS38), SBS9 (somatic hypermutation characteristic of B cells), and SBS18 (oxidative damage typically induced by reactive oxygen species). This signature decomposition was consistent with known cell type composition which included melanoma and bone marrow-derived cells. Notably, while SBS18 is not typically present in melanoma or lymphocytes^34^, it is known to occur during cell culture^35^, possibly without a subsequent clonal expansion, indicating that duplex sequencing is revealing mutational processes irrespectively of clonal expansion.

Furthermore, to assess assay-specific variation, COSMIC signature fitting was performed on COLO829-BLT50 mutational profiles generated by multiple sequencing assays (**Figure 3D**). While minor differences were observed in relative contributions of the signatures, all sequencing methods consistently recovered the core biological signal. When accounting for trinucleotide bias in each assay against WGS (**Figure S2**, **Figure S7A**), we found that these assay-specific differences had minimal impact on the COLO829-BLT50 mutational profile, likely due to limited overlap between the affected trinucleotides and the dominant features of the sample’s mutational spectrum (**Figure S7B**).

### Mutations from duplex sequencing are largely distinct from clonal reference mutations derived from bulk WGS

In addition to privately acquiring mutations, cells will also share clonal mutations due to early developmental events or somatic expansions^36,37^. Deep, non-duplex, bulk whole-genome sequencing (bWGS) has traditionally been used to detect clonal mutations and estimate their VAFs. Although duplex sequencing can detect clonal mutations^18^, it is unclear how much would duplex mutation callsets overlap with clonal callsets from bWGS on the same tissue. A small overlap would suggest that duplex sequencing could significantly expand the set of somatic mutations from bWGS analyses. Leveraging our extensive clonal “truth sets” derived from deep (∼1000X) bWGS of the COLO829-BLT50 mixture and benchmark tissues^38^, we examined its overlap with the collective sample-matched duplex mutation callsets. Simulations suggested that, on average, one unit of duplex coverage (1dX) genome-wide should recover a small fraction of reference mutations based on its size and variant allele fraction (VAF) distribution (**Figure S8A**). Aggregating across all duplex assays, we predicted that ∼11% of the reference mutations from COLO829-BLT50 (n=36,649 at VAFs of 1-2%; **Figure S8A, S8B**), <5% of reference mutations from primary tissue homogenates derived from ST002, and <10% of primary tissue homogenates derived from ST001 (∼300-500 clonal reference mutations at a broad range of VAFs; **Figure S8C**) will overlap with duplex mutation calls.

Within our data, we found that mutations identified by shallow duplex sequencing were largely distinct from the truth set (**Figure 4A**). No more than 10% of the truth-set SNVs from any tissue homogenate overlapped with its corresponding aggregated duplex callset. Except for the ST002-Lung sample, no more than 5% of truth-set SNVs overlapped with duplex calls, in line with our expectations (**Figure 4A**). Similarly, approximately 12% of COLO829-BLT50 truth set calls overlapped with duplex calls as expected (**Figure S4A**), as did the observed number of overlaps (∼4300 SNVs; **Figure S8B**). Moreover, although the aggregated duplex sequencing calls for each benchmark tissue ranged in the thousands (∼2,000-25,000), < 0.5% of those calls overlapped with the corresponding reference set. Additionally, 9-10% of the aggregated duplex sequencing calls for BLT50 overlapped with its truth set. Despite nearly all of its sSNVs occurring at 1-2%, the greater size of the BLT50 reference set likely contributed to its greater overlap with duplex calls. Despite the much smaller overlap between duplex callsets and truth sets for benchmark primary tissues, we could identify duplex mutations shared across different tissues of the same donor and single cells (**Figure S9**). Overall, the significantly lower percentages of duplex calls that belonged to the clonal reference set suggest that even shallow duplex sequencing is likely to yield a very distinct set of mutations from what can be obtained from standard deep WGS, as intuitively expected.

**Figure 4:**
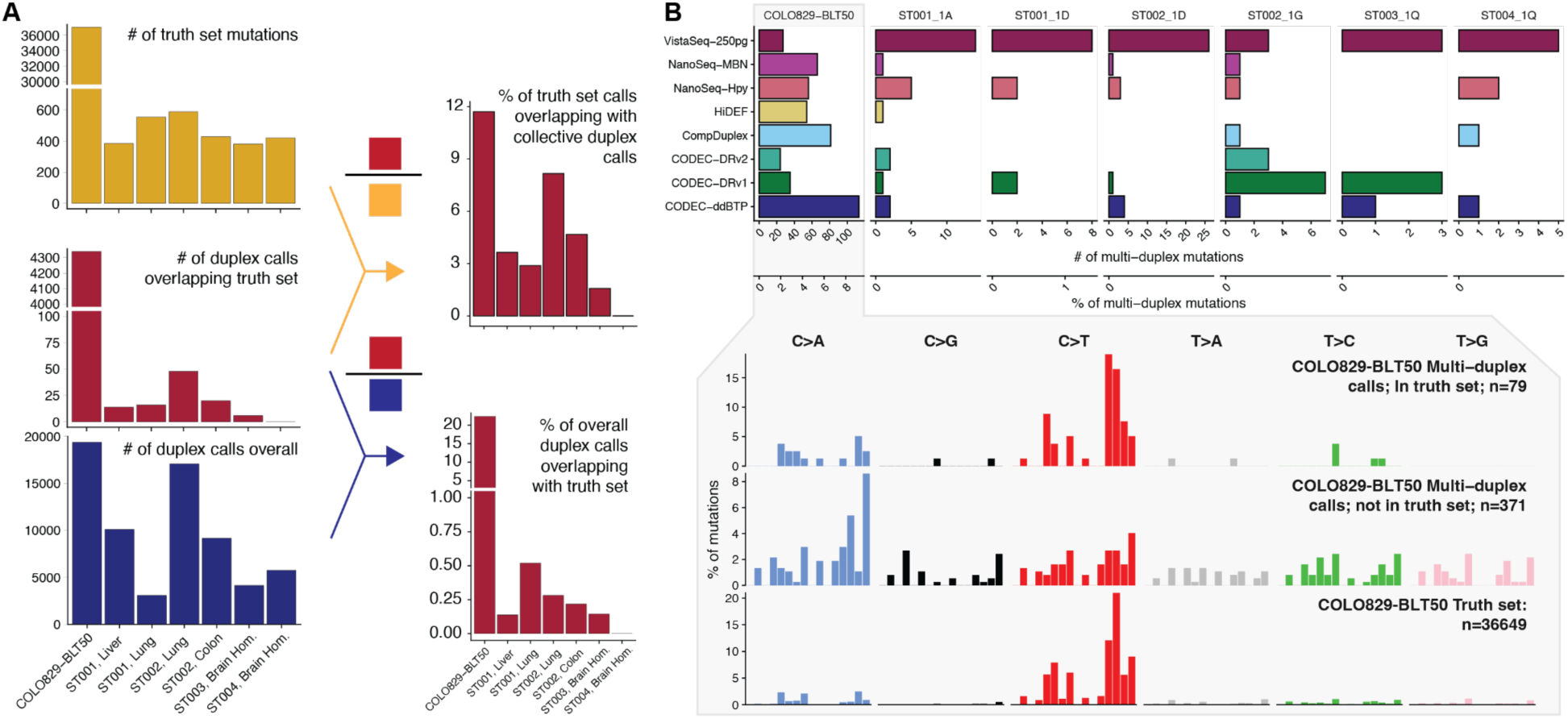
Comparison of clonal reference truth set mutations from deep bulk WGS with duplex sequencing A). Overlap of truth set mutations defined from bulk WGS and duplex mutations aggregated from different benchmarked duplex technologies. Left panel: the number of mutations in the truth set, aggregated duplex callsets, and the overlap between the two. Right panel: the percent of truth set mutations (top) or aggregated duplex calls (bottom) that are within the overlap. B). Properties of multi-duplex mutations in benchmark tissue. Top panel: the number of multi-duplex mutations identified by each technology in each benchmark tissue. The inset shows the spectra of all multi-duplex mutations identified in the BLT50 benchmark sample, separated by overlap with the truth set (middle), compared against the spectrum of the full truth set.

Multiple duplex molecules supporting a mutation would verify that it is clonal irrespective of whether matched bWGS exists. Under the infinite sites model which assumes a negligible rate of recurrent mutation at a locus^39^, a mutation in multiple passing duplex molecules implies its presence in multiple clonally-related cells’ genomic copies. We identified 450 multi-molecule mutations across all duplex sequencing assays from BLT50, 17.56% of which (n=79) also overlapped the corresponding truth set (**Figure 4B**). These overlapping mutations had a similar trinucleotide spectrum as somatic mutations called in the reference. Strikingly, the remaining 82.44% of multi-duplex mutations (n=371) formed a spectrum distinct from the truth set and devoid of its distinctive, SBS7a/b-like C>T patterns. Instead, the non-truth set multi-duplex mutations contained prominent C>A and T>G components. The contrasting spectra of two different sets of duplex-identified clonal mutations (either overlapping the truth set or not) provided further support that even shallow duplex sequencing captured very distinctive clonal populations not within the reference, motivating us to conduct more granular analyses.

### Diverse mutational signatures in duplex sequencing mirror the clonal structure of the measured tissue

Focusing on the BLT50 duplex assays and moving beyond the truth set, we examined the VAF of duplex mutations as evaluated through read pileups in bWGS (∼1000X) of the BLT50 sample and its separate tumor and blood components, all of which come from the same COLO829 donor (**Figure 5A, Figure S10A, Methods**). We also incorporated the callsets from ppmSeq, which enables much deeper duplex sequencing to yield multi-duplex coverage per base and thus permits more extensive for clonal analysis. We detected 74.2% of BLT50 duplex mutations in bWGS of BLT50 and clustered them into seven distinct clonal populations (**Figure S10**). All technologies represented the seven different inferred BLT50 clonal subpopulations at relatively even proportions (**Figure 5B**).

**Figure 5:**
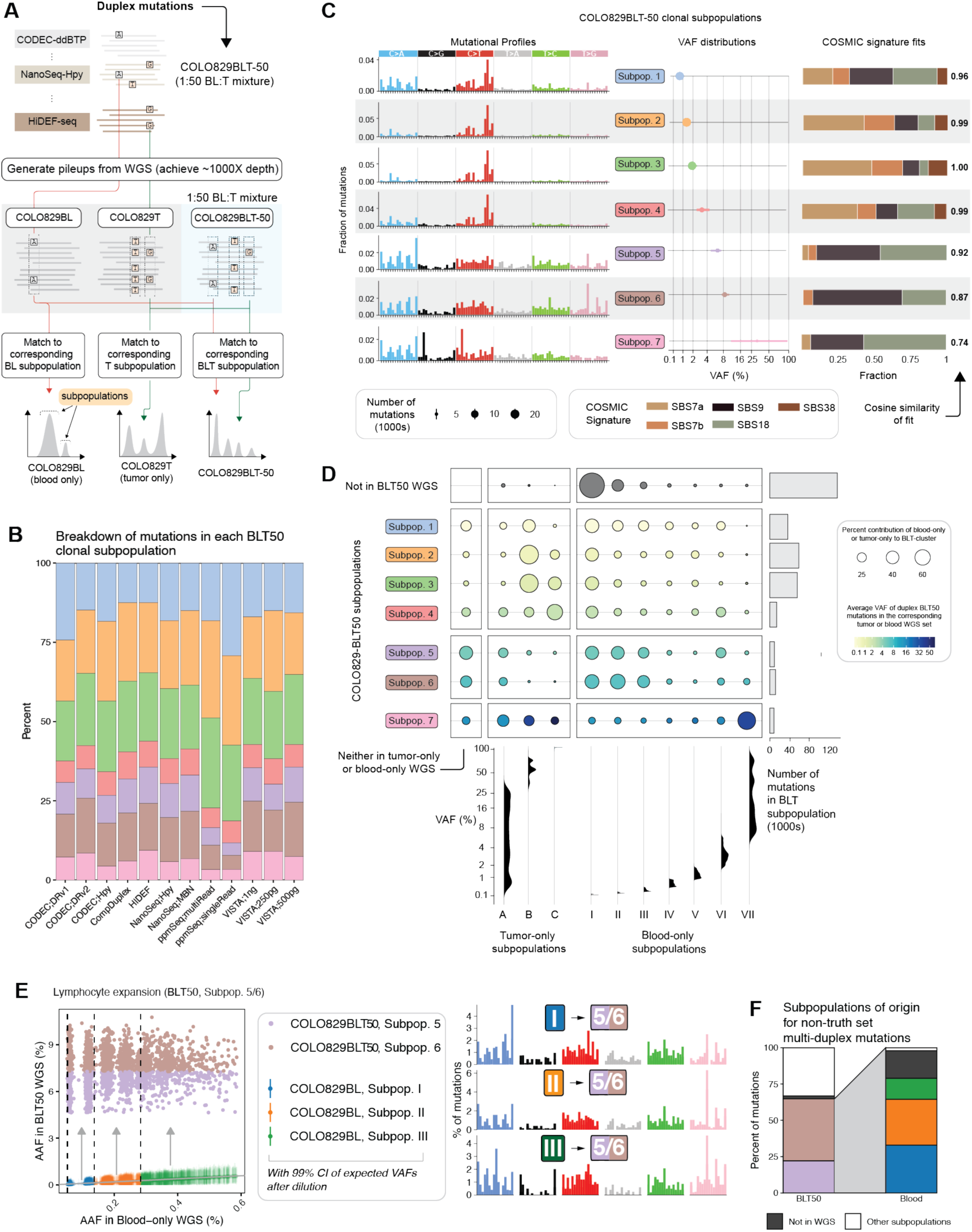
Identification of clonal populations in COLO829 BLT50 benchmark sample within duplex sequencing. A). Schematic of the expanded analysis to study the clonality of duplex sequencing-derived mutations. B). Percent of BLT50 duplex mutations assigned to each inferred BLT50 subpopulations with the procedure in (A), as broken down by technology. The trinucleotide spectrum of each inferred BLT50 subpopulation is provided. C). COSMIC mutational signatures associated with each blood, tumor, or BLT50 clonal subpopulation. D). Inferred contributions of different blood and tumor subpopulations to BLT50 subpopulations. The BLT50 sample was formed from the mixture of COLO829 blood and tumor cell cultures (at a ratio of 49:1); subpopulations of BLT50 duplex mutations are inferred to have arisen from distinct subpopulations present in the original blood and tumor cultures, all of which were determined with the procedure in (A). For each BLT50 subpopulation, the contributions of mutations coming from a tumor or blood subpopulation (whose VAFs in their respective bulk WGS samples are given in the density plots at bottom) is shown as a bubble, sized by the percent contribution of the tumor or blood subpopulation (and normalized within each row) and shaded by the VAF in the bulk WGS of BLT50. The barchart at far right shows the number of duplex mutations within each BLT50 subpopulationE). Dynamics of the lymphocyte expansion contributing to SBS9/18-associated mutations inferred from signature and clonal analysis. F). For multi-duplex mutations in BLT50 that do not overlap BLT50’s truth set (Figure 5B), shown are the percentages that belong to BLT50 subpopulations 5 and 6. The percentage that belong to other BLT50 subpopulations or not at all captured in BLT50’s bulk WGS are also shown. The percentages of multi-duplex, non-truth-set mutations in BLT50 5/6 are further broken down by the blood subpopulation. The color scheme is the same as in (E).

The trinucleotide spectra and COSMIC signature fits of the seven BLT50 clonal subpopulations (“Subpop.” 1–7) illuminated the clonal composition of the BLT50 duplex callsets (**Figure 5C**). First, large fractions of mutations in subpopulations 1–4, present at VAFs <4%, were fitted to COSMIC signatures SBS7a/b and, distinctly, SBS38. All three COSMIC signatures primarily correspond to the mutations within the tumor component of BLT50. Subpopulation 1 displayed a stronger fit to SBS18 relative to 2–4, possibly due to low-mosaic mutations formed by reactive oxygen species damage from tumor, blood, or cell culturing^40^. Subpopulations 2 and 3, in particular, occupy the VAF range that truth set mutations in BLT50 are expected to occupy (1-2% after diluting the 50-100% VAF tumor mutations in blood at a fraction of 1:50), which along with their signatures suggest that they represent the source of truth set mutations. Collectively, 51.8% of all BLT50 duplex mutations (and 84.9% of duplex mutations detected in BLT50 bWGS) arose from these four subpopulations. Second, subpopulations 5 and 6 were devoid of the SBS7a/b peaks but instead contained much more prominent C>A and T>G components, likely leading to the strong fits of SBS9 and SBS18. As the blood component of BLT50 contains lymphocytes, and as subpopulations 5 and 6 are present at VAFs of 6-8%, we hypothesize that 5 and 6 may comprise large clonal subpopulations of lymphocytes from the blood component. Finally, subpopulation 7 more closely resembled 5 and 6 than 1–4 in COSMIC fits. Subpopulation 7 fits well to SBS18, but it is devoid of both the T>G peaks in 5 and 6 and the SBS7a/b peaks in 1–4. Moreover, it shows a relatively lower cosine similarity to the spectrum constructed from its COSMIC fits compared to the other 6 subpopulations. Thus, we did not analyze subpopulation 7 further.

### Integration of duplex sequencing and ultra-deep bulk sequencing of BLT50 effectively captures a broad range of distinct clonal populations from tumor and blood components

To dissect the contributions of blood and tumor subpopulations to those seen in BLT50, we traced each BLT50 duplex mutation to one of seven blood subpopulations or three tumor populations, each of which was inferred using VAFs from pileups of BLT50 duplex mutations in blood and tumor bWGS (**Methods, Figure S10B**). We measured the number of BLT50 duplex mutations shared between each BLT50 inferred subpopulation and each blood- or tumor-only inferred subpopulation (**Figure 5D**). We noted three major trends in the correspondence of BLT50 subpopulations with blood and tumor subpopulations.

First, mutations BLT50 subpopulations 1–4 drew most of their contributions from two tumor subpopulations, “B” and “C”. Subpopulations B and C are present at average VAFs of ∼50% and ∼100% (**Figure S10C**) in the tumor bWGS as expected for truth-set mutations and which are rich in SBS7a/b and SBS38 (**Figure S10D**), respectively. Tumor subpopulation B comprised not only mutations at bWGS VAFs of 50% but also at mutations at ∼33% or ∼66% VAFs (**Figure S10C, 5D**), likely corresponding to copy number variants within the tumor component skewing the VAFs.

Second, duplex sequencing detected rare blood- and tumor-contributed mutations that could not be found in BLT50’s bWGS, hinting at the clonal resolution of duplex callsets. Duplex sequencing of BLT50 detected rare blood and tumor-contributed mutations, from blood subpopulation I-III at VAFs 0.1-1% and tumor subpopulation A at VAFs 0.1-30%, that did not appear within BLT50’s bWGS (**Figure 5D**). Although A’s mutations comprise <5% of the BLT50 duplex mutations not detectable in BLT50 bWGS, the 1:50 dilution of tumor in BLT50 implies that A would likely occur at final VAFs of 0.002-0.6% in BLT50. Moreover, I-II’s mutations, which make up the vast majority of BLT50 duplex mutations that could not be found in BLT50’s bWGS, occur mostly at 0.1-0.6%. BLT50 duplex mutations that could not be found in BLT50 bWGS make up 25.8% of all BLT50 duplex mutations, suggesting that duplex sequencing can robustly detect mutations at clonal VAFs below 0.6% (3 in 5000 cells) and as low as 0.002% (approximately 2 in 100,000 cells), far exceeding the resolution of bWGS.

Third, BLT50 subpopulations 5 and 6, which are rich in SBS9/18, receive significant contributions from blood subpopulations (I-III) but minor contributions from the tumor (**Figure 5D**). Unusually, the VAFs of these mutations seem to be greater in BLT50 than in blood, which is subject to a 49/50 dilution to create BLT50 and thus should see its subpopulations largely retain their VAFs. However, blood subpopulations I-III occur at VAFs <1% in blood bWGS but appear at VAFs of 6-8% in BLT50’s bWGS, suggestive of a clonal expansion. Overall, integrating duplex sequencing and bWGS enable a high-resolution analysis of the contributions of blood and tumor subpopulations to expected and seemingly emergent populations in BLT50.

### Integrative analysis of duplex sequencing and bulk WGS further tracks an acquired lymphocyte expansion in BLT50

We further focused on BLT50 subpopulations 5 and 6 (“5/6”), which we hypothesized represents a clonal lymphocyte expansion that occurred during the mixture of tumor and blood into BLT50^38^. Indeed, SBS9 has been associated with polymerase eta mutations made as part of somatic hypermutation during B-cell expansions ^34^. The speed and scale of lymphocyte expansions could also explain how a blood population at 0.1% VAF suddenly emerges at 8% VAF in a short timeframe. Mutations shared between subpopulations 5/6 and progenitors in blood, mostly subpopulations I–III (**Figure 5D, 5E**), displayed a significantly higher VAF in BLT50 bWGS than expected after the dilution of the blood mutations’ VAFs. In the absence of clonal expansion, the VAFs of mutations from I–III would shift from 0.1% in the blood’s bWGS to approximately 0.098% in BLT50’s bWGS (as the blood component contributes 49/50 of BLT50’s cellular content) (**Figure 5C**). The trinucleotide spectra of mutations in each of I–III contributing to 5/6 showed a consistent T>G- and C>A-rich profile consistent with SBS9 and SBS18, respectively. Although SBS18 may emerge due to cell culture^41^, it can co-occur with SBS9 in memory B-cell activation and expansion^34^. Thus, our duplex sequencing and bWGS analysis tracks the progenitors, scale, and mutational signatures of an unexpected, acquired lymphocyte expansion during the creation of BLT50.

The I/II/III-to-5/6 spectra closely resembled that of the multi-duplex mutations that did not overlap the BLT50 truth set (**Figure 5B**). As multi-duplex support would be sufficient to establish that a duplex mutation is clonal, we investigated if multi-duplex mutations would also be sufficient to detect the lymphocyte expansion. Indeed, we found that 64.8% of these multi-duplex, non-truth set mutations corresponded to 5/6 (**Figure 5F**), compared to only 8.8% from BLT50 subpopulations 1–4 (derived from the tumor-originating truth set mutations). In contrast, only 3.8% of multi-duplex mutations overlapping the truth set belong to 5/6 versus 96.2% from 1–4 (**Figure S10E**), in agreement with the near-identical spectrum of multi-duplex, truth-set mutations to the truth-set signature (**Figure 5B**). Of the multi-duplex, non-truth set mutations belonging to 5/6, 79% could be traced to blood subpopulations I-III (**Figure 5F**). Overall, we can conclude that most non-truth-set multi-duplex mutations from BLT50, which are verifiably clonal without the need for bWGS, can capture a new clonal expansion from lymphocytes in BLT50’s bWGS.

Taken together, our analysis of the clonal VAFs of duplex-captured mutations reveals three key insights. First, duplex mutations represent a largely distinct set from the bWGS mutations, representing a rich landscape of clonal populations. The low overlap with established truth sets (<10%; **Figure 4A**) suggests that duplex sequencing could dramatically expand upon the repertoire of clonal mutations. Integrative analysis with bWGS accurately estimated the clonal VAFs of mutations; deep targeted sequencing of specific mutations could also enable similar VAF estimation. Second, the expanded set of mutational signatures from duplex sequencing (**Figure 3**) may arise from the diverse clonal populations whose mutations can be effectively captured by duplex sequencing. In particular, oxidative damage processes and an acute lymphocyte expansion could be traced to specific clonal populations in BLT50 (**Figure 5E, 5F**), and further comparisons with blood or tumor bWGS could delineate the formation of these populations. The detection of a lymphocyte expansion is particularly striking, given the agreement of different technologies on the VAF and abundance of the clones relative to others. Finally, our analysis suggests that duplex sequencing has a significantly expanded sensitivity for ultra-low VAF clones well below 1% cell fraction, currently the limit for most deep bWGS (10^2^X of coverage). Duplex technologies could reliably capture mutations present at VAFs of 0.1%, with a minority of mutations appearing as low as 10^-3^-10^-5^. Although deep sequencing would be required to verify the VAFs of mutations, shallow duplex sequencing is sufficient to capture distinct signatures of low-VAF clonal populations, pointing to the ability to profile clonal expansions using lighter sequencing requirements than deep bWGS. Overall, the results support the ability of shallow duplex sequencing to study somatic mosaicism at high resolution and foreshadow potential clinical applications to detect clonal expansions in human tissue from rare progenitors^23,42^.

### Duplex consensus efficiency and economic performance

Finally, we quantified the efficiency with which raw reads are converted to high-fidelity duplex consensus sequences using two metrics: (i) duplex reconstruction efficiency, the fraction of input molecules that pass all filters to yield duplex-validated molecules (**Figure. 6A**, **Table 1**); and (ii) cost efficacy, the number of interrogated base pairs after all filters are applied (i.e., the effective depth) per unit library cost, explicitly accounting for sequencing platforms (**Figure. 6B**).

**Figure 6.**
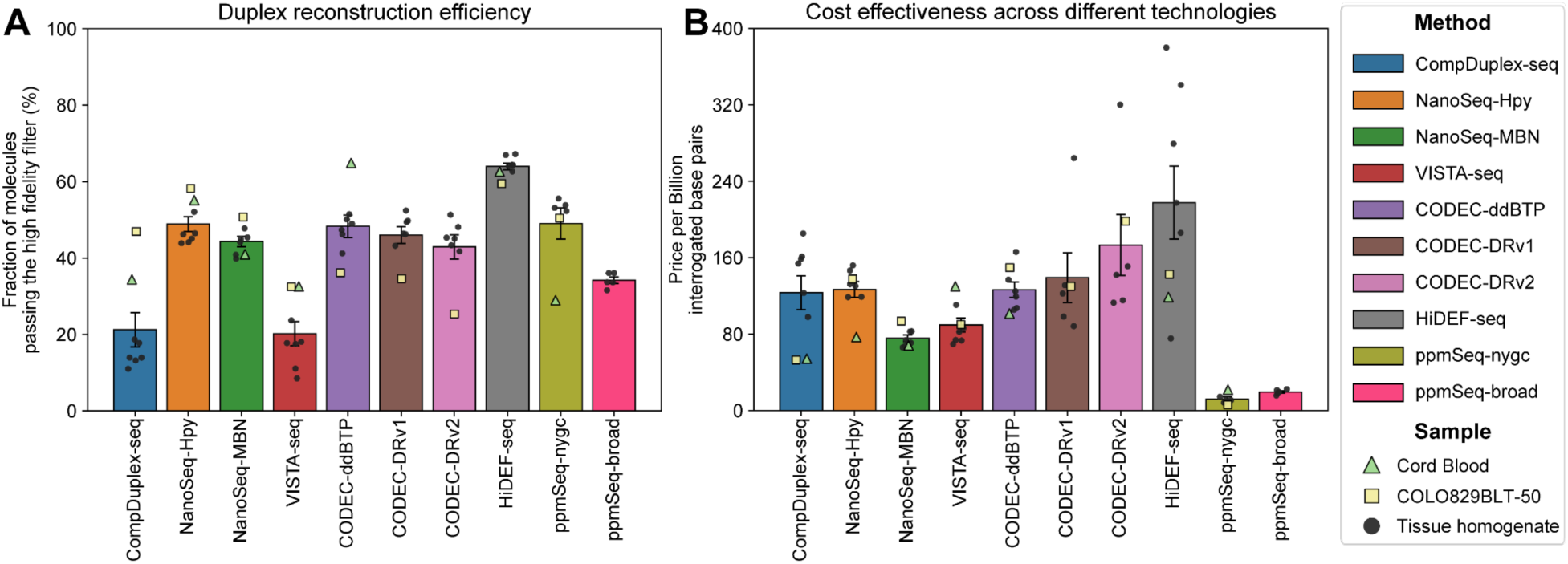
Duplex reconstruction efficiency and cost-effectiveness. A. Duplex reconstruction efficiency, defined as the fraction of unique molecules that yield a duplex consensus (both strands observed) among all sequenced molecules. B. Cost-efficacy, expressed as USD per billion interrogated base pairs. Values are averaged across all benchmarked samples. Pricing assumptions: Illumina NovaSeq X, $2/Gb (applied to NanoSeq, CompDuplex-seq, VISTA-seq); Illumina NovaSeq 6000, $5.6/Gb (CODEC); PacBio Revio, $995/SMRT Cell (HiDEF-seq; converted to $/Gbp using observed output); Ultima UG100, $1/Gb (ppmSeq). Error bars represent mean ± s.d. across samples.

In strand-barcoded workflows using high-integrity DNA samples, NanoSeq and CompDuplex-seq achieve mean duplex-recovery rates of ∼50–60%, whereas VISTA-seq averages 30.7%, reflecting its lower strand-tag extension efficiency and larger number of PCR cycles. Although strand-barcoded libraries typically require a higher number of reads per molecule duplication (≈6–8 read pairs per molecule), their high recovery and broadly usable read lengths render cost effectiveness comparable to physically linked strategies. CompDuplex-seq shows a bimodal cost profile driven by DNA integrity: it is highly efficient in high-quality cell-line DNA ($50 per billion interrogated base pairs) but incurs strand-dropout costs in tissue homogenates with elevated nicking ($150 per billion interrogated base pairs). By contrast, VISTA-seq and NanoSeq-MBN deliver more stable recovery across samples and, on average, above-median cost effectiveness (VISTA-seq: $96.1; NanoSeq-MBN: $76.24 per billion interrogated base pairs).

Physically linked duplex platforms show wider spread in cost effectiveness because each targets a distinct instrument and read architecture. On Illumina, CODEC links complementary strands via an adaptor-quadruplex so both strands are captured in a single read pair. Across CODEC designs, productive duplex recovery ranges from ∼25–36%. By definition, only the strand-overlap is interrogated for variant calling; consequently, cost effectiveness depends on insert size and read length. With appropriately long inserts/reads, high-fidelity overlap can approach the cost profile of strand-barcoded methods (≈$126 per billion interrogated base pairs). Notably, the two whole-genome CODEC libraries (DRv1/2) from one brain tissue were outliers, due to poor library qualities. HiDEF-seq on the PacBio Revio instrument captures most productively-sequenced molecules as duplexes that typically span 3–5 kb, though after filtering, ∼65% of molecules remain. Although PacBio long-read sequencing carries a higher per-base price, HiDEF-seq achieves relatively high per-read duplex yield, mitigating its cost to ≈$220 per billion interrogated base pairs. Finally, ppmSeq on the Ultima UG100 uses cluster-level, two-strand consensus emitted as a single read to exploit very high throughput. In our benchmark, its cost efficacy, at approximately $15.4 per billion interrogated base pairs, reflects the interplay between platform pricing and consensus yield rather than library construction overhead (**Figure 2D**). Taken together, duplex-recovery and cost-normalized effective depth expose clear trade-offs between molecular yield and spend, clarifying when each platform offers the most economical route to high-confidence somatic variant detection.

## Discussion

Duplex sequencing has revolutionized the study of somatic mutation patterns by enabling detection of ultra-rare variants with unprecedented fidelity. Here, through a comparison of a suite of these duplex sequencing methods on coordinated benchmarking samples, we provide a head-to-head comparison of these novel methods. The different duplex sequencing methods were in broad agreement on estimated mutation burden and mutational signatures, highlighting the ability of all of them to distill biologically consistent insights from the data. Nevertheless, differences in chemistry and sequencing platforms yield differences in genomic footprints, cost efficiencies, and subtle differences in error profiles and trinucleotide abundances.

Comparative analysis of duplex sequencing with ultra-deep matched bulk sequencing revealed the ability of duplex sequencing to interrogate somatic mutation patterns arising across the entire lifespan. Consequently, duplex sequencing yields mutations largely distinct from bulk sequencing. Mutation calls from bulk sequencing appear confined to early events or highly clonally expanded mutations, whereas duplex sequencing permits the detection and inference of late, often tissue-specific mutational processes, for example, smoking in lung and a liver-specific mutational signature. Our observations on mutational burdens and signatures are consistent with single-cell DNA sequencing data from the SMaHT Network^9^. One advantage of duplex sequencing is its ability to measure average mutation burdens and signatures within a population of cells from a single sample, whereas single-cell DNA sequencing has to rely on sampling a sufficient number of single cells to generate a representative picture of mutagenesis within a tissue.

Duplex sequencing is not without limitations. Currently, duplex error correction works well for short mutations whose span lies within sequencing reads, but it is unable to precisely detect copy number variants or structural variants, likely due to an even lower rate of somatic structural variation than SNVs. Background processes in duplex sequencing generating chimeric reads currently overwhelm the rate of true somatic structural variants. In parallel, the error rate of short insertions and deletions (indels) is highly dependent on sequencing chemistry, and currently only the Illumina-based methods work for indel detection.

The choice of duplex sequencing approach for a given experiment will largely be driven by the biological questions, sample requirements, and amount of input material. For example, HiDEF-seq can deconvolve single-strand mismatches, damage, and modifications concurrently with double-strand mutations^20^. VISTA-seq works well on variable amounts of input DNA, even as low as a single cell, for both SNVs and indels^43,44^. NanoSeq has been used extensively in the past years, generating genome-wide mutation patterns and, in combination with bait capture, precise signals of genes under selection in normal tissues^23,24^. CompDuplex enables single-cell duplex-seq profiling and more importantly integration with a total-RNA based single-cell transcriptome assay^45,46^ for single-nucleus dual-omics profiling of frozen tissues. CODEC has been applied and tested well on cell-free DNA for liquid biopsy studies with low amounts of DNA^18^ and can be paired with methylation inference^47^. PpmSeq natively supports duplex reconstruction on the Ultima Genomics sequencing machine and can generate ultra-deep genome-wide duplex data, currently employed for detection of minimal residual disease in liquid biopsies^21^.

The development and application of duplex sequencing is a highly active area in genomics research. Beyond the methods tested in this study as part of the SMaHT Network, ultradeep targeted duplex sequencing is used to detect signals of selection^42^, new variations of duplex sequencing continue to improve coverage and efficiency^48^, and the bioinformatics around duplex sequencing will be further developed ^49^. In parallel, further reductions in sequencing cost may eventually overcome the trade-off between genomic coverage and depth and to bridge single-molecule and clonal mutational landscapes. At present, the various methods of duplex sequencing offer researchers a unique and adaptable toolkit to study mutational landscapes across tissues, discover driver mutations, and detect minimal residual disease in cancer patients. We foresee that the lessons learned from this benchmarking, and the integrative analyses, are widely applicable both within the production phase of the SMaHT Network and in the wider somatic mutation field

## Competing interest declaration

V.A.A. and R.L. are co-inventors on patent applications filed by the Broad Institute on CODEC and Methyl-CODEC. V.A.A. is a co-inventor on patent applications filed by the Broad Institute on Duplex-Repair and MEDUSA, and on the MAESTRO MRD test which has been licensed to Exact Sciences. V.A.A. is a co-founder and advisor to Amplifyer Bio. G.D.E. has filed a patent application for HiDEF-seq and owns equity in DNA sequencing companies (Illumina, Oxford Nanopore Technologies, and Pacific Biosciences). C.Z. is a co-founder and equity holder of Pioneer Genomics and reports that Baylor College of Medicine filed a patent application related to the CompDuplex-seq or CompDup method. R.A.G and Baylor College of Medicine have equity interest in Codified Genomics, Ltd.

## Acknowledgements

We are extremely grateful to the SMaHT donors, and donor families, who have generously provided such precious gifts to support this important work. This research is supported by the NIH Common Fund, through the Office of Strategic Coordination/Office of the NIH Director under awards U24 MH133204, U24 NS132103, UG3 NS132132, UG3 NS132024, UG3 NS132144, UM1 DA058229, UM1 DA058230, UM1 DA058235, and UM1 DA058236. The NYGC and the Broad Institute have received material support from Ultima Genomics for data generation for this study.

## Author Contributions

**Writing**: Y.Z., V.V.V., M.A., D.G., G.D.E., C.Z. and T.H.H.C., with input from all other authors Figures: Y.Z., V.V.V., M.A.

**Variant calling**: R.L. and N.L. (CODEC); Y.Z. (NanoSeq and CompDuplex-seq); S.-Y.J., G.D. and M.D.M. (VISTA-seq); M.G.-P. and G.D.E. (HiDEF-seq); L.J.L. and U.S.E. (ppmSeq)

**Analyses**: Y.Z. and M.N. (metrics and burdens); M.A., D.G., H.J., C.S. and T.H.H.C. (mutational signatures); V.V.V. and J.A.B. (clonal analyses)

**Experimental work**: A.N., M.N., H.C., A.H., N.T.J., S.L.M., C.S., N.T.J., N.H.

**Data and project support**: L.A., C.C. W.C.F., A.R.

**Study design**: K.G.A., S.G., R.A.G., S.C., H.V.D., G.D.E., C.Z. and T.H.H.C.

**Supervision**: D.S., C.A.W., V.A.A., E.A.L., P.J.P., K.G.A., S.G., R.A.G., S.C., H.V.D., G.D.E., C.Z. and T.H.H.C.

## Data Availability

The datasets described in this study will be made available through dbGaP under the study accession numbers phs004193. The data used in this work was provided by the SMaHT Data Analysis Center (DAC) on behalf of the SMaHT Network. More information about the SMaHT Network and data is available online at https://smaht.org and at https://data.smaht.org. Data reuse should abide by the SMaHT data policy (https://smaht.org/data-use-policy/).

## Methods

### 1. Library preparation and sequencing

**ppmSeq:** ppmSeq (Paired Plus Minus Sequencing) libraries were prepared using the NEBNext UltraShear module (NEB M7634L) and NEBNext Ultra II DNA PCR-free Library preparation Kit (NEB E7410L) in accordance with the manufacturer’s instructions. 250ng of DNA was sheared enzymatically and was subsequently end-repaired and adenylated. DNA fragments were ligated to Ultima Genomics Indexed Native Duplex Adapters and the libraries underwent bead-based size selection. Final libraries were quantified using the QuantStudio5 Real-Time PCR System (Applied Biosystems) and Fragment Analyzer (Agilent). ppmSeq libraries were sequenced on Ultima Genomics (UG100) instruments according to manufacturer instructions.

**NanoSeq:** NanoSeq HpyCH4V and NanoSeq Mung Bean libraries were prepared using 50-100ng of DNA as described previously^17^ with minor changes as described in Chao et al.: https://www.protocols.io/view/optimized-mung-bean-nuclease-nanoseq-library-prepa-d4h38t8n.html.

NanoSeq libraries were pooled and sequenced on Illumina NovaSeq sequencing platforms for 30X coverage. Analysis details are described in the Duplex sequencing companion manuscript.

**CompDuplex-seq:** CompDuplex-seq libraries were prepared using 20-50ng of gDNA as described by Niu and Zong (https://www.protocols.io/view/compduplex-accurate-detection-of-somatic-mutations-kxygx3x4og8j/v1). Libraries were pooled and sequenced on Illumina NovaSeq sequencing platforms for 30X coverage.

**CODEC:** CODEC sequencing was performed as described previously^18^. Two distinct end repair and dA-tailing strategies were implemented to suppress error propagation during library preparation as described below. ddBTP-CODEC: Genomic DNA was enzymatically fragmented with HpyCH4V and AluI (NEB) and size selected for 150bp size. 20 ng fragmented DNA was dA-tailed using Klenow fragment with a dATP/ddBTP mix (ddTTP, ddCTP, ddGTP), following the method of Abascal et al. Duplex Repair-CODEC (DRv1): Genomic DNA was mechanically sheared via Covaris to ∼150 bp, and 20 ng was subjected to end repair and dA-tailing as described by Xiong K et al. (NAR, 2022). An enhanced version of this protocol (DRv2, manuscript in preparation) was also used to further minimize damage-related artifacts. Following ER/dA tailing, CODEC specific quadruplex adapters were ligated. Libraries were PCR-amplified and sequenced on the Illumina NovaSeq 6000 (S4 flow cells).

**HiDEF-seq**: DNA of ST001-Lung, ST002-Colon, ST002-Lung, ST003-Cortex, and ST004-Cortex were extracted with the PacBio Nanobind PanDNA kit per the manufacturer’s protocol for animal tissues, except with Proteinase K and RNase A incubations at 37 °C. DNA of ST001-Liver was extracted with the Qiagen Puregene Kit per the manufacturer’s protocol for tissues, except with Proteinase K and RNase A incubations at room temperature. DNA of ST001-Liver was additionally cleaned with the Qiagen DNeasy PowerClean Pro Kit to remove inhibitors that interfere with sequencing. DNA of umbilical cord blood (StemCell Technologies, cat. # 70007.1) and COLO829BLT50 were extracted with the Qiagen Puregene Kit per the manufacturer’s protocol for cultured cells, except with RNase A incubation at room temperature.

We developed a new version of HiDEF-seq library preparation (v3) that improves on the prior HiDEF-seq protocol^20^ by simplifying the workflow and by utilizing random fragmentation for genome-wide coverage. A detailed step-by-step protocol is available^50^. We used the non-A tailing, blunt adapter ligation version of the protocol for this study’s samples to avoid single-strand mismatch artifacts that can occur due to A-tailing in the setting of fragmented post-mortem tissues^20^.

Briefly, HiDEF-seq v3 libraries were prepared by first removing short DNA fragments using the Short Read Eliminator (SRE) kit (PacBio; cat. #102-208-300) for all samples except ST001-Liver whose DNA was not compatible with SRE due to greater post-mortem fragmentation. For ST001-Liver, we instead removed short fragments with a 0.75X bead to sample volume ratio of 75% diluted AMPure PB beads (PacBio). DNA was randomly fragmented to a peak size of ∼4 kb using a Megaruptor 3 (Diagenode) fitted with short hydropores (cat. # E07010001) in a volume of 100 µL with 3 consecutive runs at speed ‘65’. DNA overhangs were blunted by incubating the DNA with Nuclease P1 (NEB) at 0.5 U/µL (buffer: NEBuffer r1.1) for 30 mins at 37 °C followed by inactivation with a final concentration of 19 mM EDTA and 0.01 % SDS and a cleanup with Ampure PB beads (PacBio) at a 1.5X bead to volume ratio. DNA was then treated with T4 PNK (NEB) at 0.4 U/µL (buffer: rCutSmart; 4mM DTT; 1mM ATP) for 60 mins at 37 °C. Next, E. Coli DNA Ligase (NEB) at 0.5 U/µL and NAD+ at 26 µM (buffer: rCutSmart) were added with incubation for 30 mins at 16 °C. Residual nicks were blocked by adding Klenow fragment 3’-5’ exo- (NEB) at 0.25 U/µL and ddBTP (without dATP) at 0.1 mM each (buffer: rCutSmart) with incubation for 30 mins at 37 °C. Adaptors were ligated in a total volume of 72.5 µL by adding pre-annealed non-T overhang hairpin adapters at a final concentration of 0.88 µM, 23 µL of NEBNext Ultra II Ligase Master Mix (NEB), and 0.75 µL of NEBNext Ligation Enhancer (NEB) with incubation for 60 mins at 20 °C. Reactions were cleaned up with a 1.2X bead to volume ratio. Non-circular molecules were removed by adding Exonuclease I (NEB) at 0.29 U/µL, RecJF (NEB) at 0.86 U/µL, and T7 Exonuclease (NEB) at 0.57 U/µL (buffer: NEBuffer 4) with incubation for 30 mins at 37 °C. Finally, libraries were cleaned up with a 0.9X Ampure PB bead to volume ratio.

Libraries were sequenced on a PacBio Revio instrument with 30 hour movies and settings: ‘Include Base Kinetics = TRUE’, ‘Sample is indexed = FALSE’, ‘Use Adaptive Loading = TRUE’, ‘Consensus Mode = strand’, ‘Full Resolution Base Qual = TRUE’, and ‘Subread To HiFi Pileup = TRUE’.

**VISTA-seq:** The VISTA-seq method was adapted from META-CS^22^ to enable the ultra-sensitive detection of SNVs and indels across a wide dynamic range of DNA input, from single cells to bulk populations, using a single-tube duplex consensus sequencing strategy (<u>Protocol.oi, 2024</u>). Building upon META-CS, which was optimized for single-cell, diploid contexts, VISTA-seq extends the framework to pooled inputs containing hundreds of alleles, where the two-allele assumption no longer applies. Core idea of this approach is the use of engineered Tn5 transposase complexes pre-loaded with strand-specific adapters, which allow for simultaneous fragmentation and tagging of both DNA strands (as in META-CS), followed by an informatic deconvolution that resolves individual molecules within multi-allelic mixtures.

A buffer containing Triton X-100, Thermolabile Proteinase K, and DTT was used for gentle lysis and protein digestion of gDNA. Tagmentation was then carried out with Tn5 transposase, which fragments DNA and incorporates barcoded adapters into both strands in a symmetric fashion.

The reaction was halted with a proteinase-containing mix, followed by strand-specific tagging through two rounds of synthesis with uniquely indexed primers (ADP1 and ADP2) and high-fidelity Q5 polymerase. Each synthesis step was succeeded by exonuclease digestion to remove unincorporated primers and reduce background. The resulting duplex-tagged libraries were PCR amplified, purified with Zymo columns, and size-selected through a two-step AMPure XP bead cleanup to enrich for specific fragment size ranges ∼400–600 bp. Key experimental differences from META-CS include the optimization for short-read sequencing 150x2bp instead of 250x2bp for cost efficiency, and modifications for further error reduction such as usage of thermolabile enzymes. Final libraries were quantified using TapeStation and sequenced on the Illumina NovaSeq X Plus 10B platform with paired-end 150 bp reads and a 20% PhiX control.

### 2. Variant calling and filtering

**ppmSeq:** ppmSeq data were aligned and preprocessed using the manufacturer’s custom pipeline and parameters except to use the hg38 human reference genome without decoy sequences to match other alignments in this study. Somatic SNVs were then called from the resulting CRAMs using a newly developed pipeline for calling SNV mutations, producing sensitivity-corrected mutational spectra and estimates of the SNV mutation rate per base pair (manuscript in preparation). This new method is applied downstream of the manufacturer’s SRSNV pipeline^21^, which produces a list of reads supporting candidate somatic SNVs (all non-reference sites) and assigns to each a quality score (the *SNVQ*, with higher values being better) encapsulating the proportion of amplicons from the Watson and Crick strands of the original DNA molecule and other important quality covariates. The input to the new SNV calling pipeline is the aligned ppmSeq CRAM, the unfiltered output of SRSNV and a single numerical parameter giving the desired false discovery rate (FDR) among the final somatic SNV calls, which was set to 10% for this study. Following the manufacturer’s recommendation, CRAM and SRSNV output outside of the Ultima high-confidence regions^51^ (∼93% of the genome) was discarded.

Briefly, SNV calling hinges on comparing the distribution of SNVQ scores at candidate somatic SNVs against the SNVQ distribution of known germline SNP variants, which is considered the distribution of true events. The number of true SNV mutations among the SNV candidate set is estimated using a procedure inspired by q-value calculation at various SNVQ cutoffs and the cutoff most closely matching the desired FDR target is chosen. Calling sensitivity is then estimated by the fraction of reads containing known germline SNVs that pass the same SNVQ cutoff. SNVQ-based calling is carried out separately for each of the 96 channels used in the SBS96 mutational signature format, allowing for corrected mutational spectra by adjusting the number of per-channel SNV calls by its sensitivity. Total extrapolated mutation burdens are the sum of the adjusted SNV call numbers per channel. The number of interrogated bases is given by the number of bases in duplex reads (MIXED-MIXED ppmSeq reads) adjusted by the loss rate of the SRSNV pipeline, estimated by the fraction of reads at homozygous SNPs in the aligned CRAM file that are retained in the SRSNV output. Mutation rates per base pair are the total extrapolated mutation burdens divided by the number of adjusted interrogated bases.

**CODEC:** CODEC data were processed as previously described^18^, except that the hg38 reference genome without the decoy sequences was used in this study. In summary, samples were analyzed using the standard CODECsuite (https://github.com/broadinstitute/CODECsuite) from BCL files to variant calls. All CODEC data was processed through a cloud pipeline based on Terra (https://terra.bio) and WDL (Workflow Description Language). The cloud pipeline is available at Dockerstore: https://dockstore.org/workflows/github.com/broadinstitute/TAG-public/SingleSampleCODEC:CODEC?tab=info.

**CompDuplex-seq and NanoSeq:** Variant calling and filtering were performed using a unified CompDuplex–NanoSeq analysis pipeline, available at https://github.com/zonglab/CompDuplex. The workflow includes adapter trimming, alignment to GRCh38, consensus molecule generation, duplex validation, and stringent variant filtering based on base quality, mapping quality, and duplex support, ensuring high-confidence detection of ultra-rare mutations.

**HiDEF-seq:** HiDEF-seq data was analyzed using a new HiDEF-seq v3.0 pipeline we developed (https://github.com/evronylab/HiDEF-seq) that is compatible with data produced with PacBio Revio instruments. The new pipeline follows the same read processing and filter steps as the previously published HiDEF-seq pipeline^20^ but has been fully refactored to utilize Nextflow workflow management, an updated docker image (docker hub: gevrony/hidef-seq:3.0), and more efficient scripts that process data in parallel chunks to reduce maximum memory requirements for each individual process.

**VISTA-seq:** VISTA-seq data were processed, and somatic SNVs were called using a customized pipeline that integrates elements from *pre-pe* (https://github.com/lh3/pre-pe) and *lianti* (https://github.com/lh3/lianti), with substantial modifications to support multi-allelic input. A key innovation was the introduction of barcode extraction and pooled read handling. During pre-processing, only reads with a complete matching barcode sequence to the reference list were retained, and barcodes were extracted without imposing strict requirements on transposon binding sequence matches—reducing read loss from PhiX-induced errors at transposon binding sites. Because each original DNA molecule was identified by a pair of barcodes, aligned reads sharing the same barcode pair were pooled prior to pileup, enabling accurate candidate calling in contexts where hundreds of molecules may be present at a single locus. This pooling step was essential for extending analysis beyond diploid assumptions and accommodating bulk inputs. In addition, we performed further filtering of recurrent false positives identified through rigorous benchmarking, including variants with extreme allelic imbalance under multi-allelic assumptions and variants located too close to read ends. These additional filters were critical for improving specificity in pooled-input contexts.

### 3. Metrics

#### All metrics can be found in Table S1. Interrogated Base-Pair Calculation

To enable fair cross-platform comparison, we defined *interrogated base pairs* (IBP) as the total number of genomic bases with high-fidelity for variant detection after duplex consensus formation and applying all filters, regardless of the underlying molecular barcoding strategy.

Each technology developer group applied its platform-specific pipeline, whether strand-barcoded, physically linked, or enzymatically repaired, to assemble duplex consensus molecules and perform variant calling. These high-fidelity molecules represent genomic fragments for which both complementary strands were independently sequenced and mutually validated. All genomic positions covered by these duplex-validated molecules and passing standard base-quality, mappability, and genome-mask filters collectively define the *effective callable genome*. Conceptually, the interrogated base pairs (IBP) quantify the total number of high-fidelity bases available for confident single-nucleotide or small-indel variant detection. Mathematically, for sample *i* with 𝑁_!_ duplex molecules:

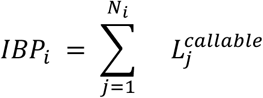

where 𝐿^&’((’)(*^is the number of callable bases within molecule *j*. This metric, expressed in base pairs, normalizes for differences in molecule yield, barcoding scheme, and consensus stringency, providing a chemistry-agnostic denominator for downstream comparisons such as mutation burden, coverage modeling, and cost-effectiveness analysis.

#### Cost-Efficacy Benchmarking

To assess the trade-off between sequencing yield, fidelity, and cost, we defined cost effectiveness as the sequencing cost per billion IBPs, normalized to the point of maximum calibrated sensitivity:

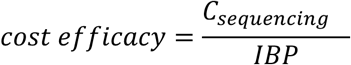

where 𝐶_+*,-*.&!./_ denotes the sequencing cost required to reach maximal calibrated sensitivity, i.e., the saturation point beyond which additional depth yields negligible increase in variant recall. For each technology, the analysis pipeline was tuned to achieve consistent accuracy on the low-mutation-burden cord blood reference while maximizing sensitivity across callable loci. The resulting *cost-effectiveness* values provide a cost-normalized measure of price per billion IBP, enabling direct comparison across platforms irrespective of chemistry complexity, barcoding architecture, or sequencing instrument. This framework quantitatively integrates experimental efficiency and economic feasibility under a unified analytical benchmark.

#### Empirical Coverage Distribution

For each library, genome-wide duplex coverage was defined as the number of distinct duplex molecules covering each locus:

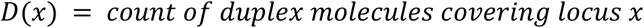

The empirical distribution 𝑃(𝐷 = 𝑑) summarizes coverage uniformity for both total and high-fidelity molecules. Under ideal random fragmentation and unbiased sampling, duplex molecules are expected to follow a *Poisson process* with mean coverage 𝜇:

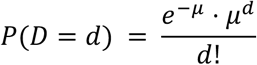

However, empirical data often deviate from this model due to preparation-specific biases. Restriction-enzyme libraries exhibit local over-dispersion caused by uneven cut-site spacing and sequence-context-driven alignment anchoring, producing clusters of molecule start sites and zero-coverage gaps. Sonication-based libraries typically show more uniform coverage, yet minor deviations persist due to stochastic ligation and mappability constraints.

#### Zero-Inflated Negative Binomial (ZINB) Modeling

To accurately capture both the excess of zero-covered loci and the over-dispersed tail of the coverage distribution, we fitted a *Zero-Inflated Negative Binomial (ZINB)* model:

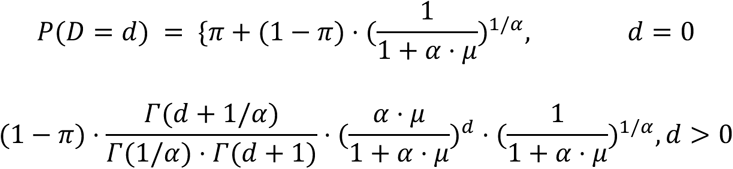

where:

● 𝜋: fraction of structural zeros representing unfragmented or inaccessible loci,
● 𝜇: mean duplex coverage,
● 𝛼: the *over-dispersion parameter*, with variance given by 𝑉𝑎𝑟(𝐷) = 𝜇 + 𝛼 ⋅ 𝜇^5^,
● 𝛤(⋅): the Gamma function.

Under this parameterization, 𝛼 → 𝟎 corresponds to the *Poisson limit* (i.e., purely random fragmentation without overdispersion), while α > 0 captures extra-Poisson variability arising from library-level biases such as sequence-context clustering, ligation preference, or alignment anchoring.

Model parameters were jointly estimated by maximum likelihood optimization implemented in a customized Python script using numerical solvers from the SciPy library (scipy.optimize). Initial parameter seeds were obtained from empirical moments of the observed coverage histogram, and convergence was verified by likelihood curvature and parameter stability.

Across all library designs, this custom ZINB fitting achieved a better fit with empirical distributions compared with the Zero-Inflated Poisson (ZIP) approximation (**Figure S1**), particularly for restriction-enzyme–based libraries where fragmentation-site density induces structured dropout. This probabilistic modeling provides a unified quantitative framework for comparing duplex coverage uniformity across distinct NanoSeq optimization strategies.

#### Trinucleotide Context Enrichment

Another factor influenced by biased genomic sampling is the *trinucleotide context enrichment* (TCE) of interrogated bases relative to the baseline genome composition, as deviations in context accessibility can distort observed substitution profiles. The reference genome (hg38, excluding masked regions) was decomposed into all contiguous 3-bp represented in a pyrimidine-centric orientation, such that each context is defined with respect to the reference C or T base at the central position (**Figure S2A**). For each trinucleotide class 𝑘 ∈ {𝐴𝐶𝐴, 𝐴𝐶𝐶, . . . , 𝑇𝑇𝑇}, baseline counts were recorded as 𝐶_/_(𝑘). The same counting procedure was then applied to the reference coordinates underlying each sample’s IBPs, yielding 𝐶_!_(𝑘). Normalized frequencies were obtained as:

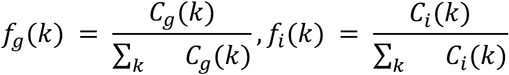

and the enrichment ratio for each context was defined as:

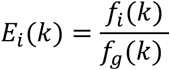

Values 𝐸_!_(𝑘) > 1 indicate relative over-representation and 𝐸_!_(𝑘) < 1 depletion of context 𝑘 within the interrogated regions. This ratio, while commonly termed “enrichment,” can equivalently be interpreted as a *context-specific accessibility ratio*, representing the relative sampling probability of each trinucleotide under a given sequencing chemistry. The resulting 32-element enrichment vector per library on COLO829BLT (**Figure S2B**) provides a compositional fingerprint for assessing systematic accessibility bias. To quantitatively evaluate trinucleotide bias, the standard deviation among normalized enrichment values was calculated and displayed in the legend of **Figure S2B**, where values approaching zero indicate minimal bias.

Next we compared technologies employing distinct fragmentation strategies, methods based on random genome fragmentation consistently demonstrated lower trinucleotide-context bias (**Figure S2C**). A pronounced depletion at the GCA trinucleotide corresponds to the loss of HpyCH4V recognition sites (T|GCA) in restriction enzyme–based workflows.

When comparing the total versus filtered profiles (see *Interrogated Base Pairs* section), in strand-barcoded methods (**Figure S2C i-iv**), the filtered profiles exhibited a slightly increased bias, presumably due to the requirement of more than one read per strand. This criterion amplifies PCR-induced coverage variation, leading to preferential sampling of fragments with distinct GC-content profiles and, consequently, slightly higher trinucleotide bias. In contrast, physically linked-read methods show divergent behavior: for CODEC (**Figure S2D v–vii**), the a2s1-filtered regions exhibit reduced trinucleotide-context bias relative to all interrogated bases, whereas for HiDEF-seq (**Figure S2D viii**), the pass regions display a more pronounced bias. The latter likely reflects trinucleotide-context–dependent error preferences of the PacBio sequencing platform, which differ from those of Illumina.

Finally, nearly all technologies exhibited reduced trinucleotide-context bias in Cord Blood compared to COLO829 (**Figure S2E**), reflecting the higher genomic integrity and diploid stability of normal tissues. In contrast, the COLO829 tumor spike-in sample likely introduces compositional distortions arising from copy-number variations (CNVs) and other tumor-specific genomic aberrations.

#### Single-strand events and random error rates

Duplex sequencing distinguishes complementary strands, enabling detection of single-strand events that occur within double-stranded regions. Among the six benchmarked duplex-sequencing technologies, four provide per-read strand assignment (via native/synthetic barcodes or orientation flags), permitting strand-resolved single-strand events calling. For each trinucleotide context 𝑘 ∈ {𝐴𝐴𝐴, . . . , 𝑇𝑇𝑇} (𝑛 = 64) and each of its three possible substitutions, we tally strand-resolved single-strand events using SAM flags and read orientation. Let:

● 𝐼𝐵𝑃(𝑘): the number of interrogated base pairs in trinucleotide context kkk; for duplex analyses this equals 2× the 96-class pyrimidine-centric IBP count because both strands are considered (see *Trinucleotide Context Enrichment*).
● 𝑁_++_(𝑘, 𝑠): observed count of single-strand events of type 𝑠 in context 𝑘.

Because opportunities exist on both strands, the per-base single-strand events potential (rate) for

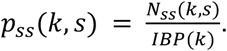

These context-specific single-strand event potentials are summarized in **Figure S3** (upper panels).

From single-strand potentials to duplex-like random errors. A false duplex variant arises when *independent* SSEs occur on both strands at the same locus in the complementary forms that mimic a true mutation. Let [inline] be the library’s trinucleotide composition. Denote by <u>𝑘</u> the reverse-complement context and by <u>𝑠</u> the complementary substitution (e.g., C>A,A-T pairs with G>T,A-T). Under independence,

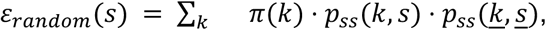

which yields the expected duplex-like random error rate for substitution class sss. We display the full profile {𝜀_?’.2@A_(𝑠)} in **Figure S3** (lower panels).

These estimates are lower bounds on the false-positive rate because detectability of damage is polymerase- and chemistry-dependent. For example, 𝐺 → 𝑇 often reflects 8-oxo-G, whereas 𝐶 → 𝑇 can arise from 5-mC deamination; if a polymerase fails to convert a damage type into a detectable misincorporation on one strand, the corresponding 𝑝_++_(𝑘, 𝑠) - and thus 𝜀_?’.2@A_(𝑠) - will be underestimated. Consequently, 𝜀_?’.2@A_(𝑠) quantifies the random-coincidence floor rather than the complete false-positive burden, which also includes systematic and context-coupled errors.

### 4. Mutational signature analysis

#### All outputs from mutational signature analysis can be found in **Tables S2-9**. **Sample selection for main signature discovery**

Prior to performing signature discovery, mutational spectra were compared across all technologies and samples and found to be largely consistent, with one exception: ppmSeq multi-duplex mutations (**Figure S5**). PpmSeq data were stratified by read support into “single” (supported by one read) and “multi” (supported by multiple reads) groups. The multi group showed consistent differences relative to other technologies in all samples, except COLO829-BLT50, which harbors a high number of clonal mutations. Signature discovery on the full catalog, including both single- and multi-duplex ppmSeq mutations, identified technology-specific signatures (Sig. 6–8; **Figure S6**), likely reflecting a residual fraction of false-positive calls given the unprecedented number of mutations identified. To minimize confounding of *de novo* signatures by potential false-positive calls, and since ppmSeq is an emerging technology, ppmSeq mutations were excluded from the main signature discovery. Because the ppmSeq mutation profiles in COLO829-BLT50 closely resembled those from other technologies, COSMIC signatures were fitted directly to this dataset (Figure 4D).

#### Signature discovery using MuSiCal

For de novo signature discovery, we used MuSiCal’s standard non-negative matrix factorization (NMF) framework with default parameters. To interpret the mutational processes captured by the de novo signatures, we decomposed each signature into a linear combination of known COSMIC v3.4 single-base substitution (SBS) signatures. To enhance biological relevance and minimize spurious assignments, the reference set of COSMIC signatures was restricted to those previously reported in the same tissue type or sample context^17,31,52–54^. Overall, the set of tissue-relevant signatures included the following COSMIC v3.4 signatures: SBS1, SBS2, SBS4, SBS5, SBS7a, SBS7b, SBS7c, SBS7d, SBS8, SBS9, SBS12, SBS13, SBS16, SBS17a, SBS17b, SBS18, SBS19, SBS22a, SBS22b, SBS28, SBS29, SBS37, SBS38, SBS39, SBS88, SBS40a, SBS89, SBS23, SBS24, SBS41, SBS84, SBS85, SBS94, SBS97.

The decomposition was performed using MuSiCal’s multinomial likelihood–based sparse non-negative least squares (NNLS) model. Contribution coefficients were estimated via maximum-likelihood refitting using a bidirectional likelihood model, with the threshold parameter set to 0.01. The cosine similarity between the reconstructed profile and the original de novo signature, reflecting the reconstruction accuracy, was reported for each signature.

#### Signature discovery using HDP

To confirm the mutational signature results from MuSiCal, we used the Hierarchical Dirichlet Process (HDP, https://github.com/nicolaroberts/hdp) for de novo signature extraction. HDP was run using default parameters (4 independent chains, a burn-in of 10,000 iterations, 200 samples with a spacing of 200 iterations). Both a tissue-based hierarchical architecture (grouping all tissue across different technologies into parent nodes) and a technology-based hierarchy (grouping all samples from one technology across tissues into parent nodes) revealed the same number and composition of extracted signatures (n=6, see **Figure S6A**). Comparison between MuSiCal and HDP signatures revealed very high concordance between the two methods (**Figure S6B**).

### 5. Analysis of clonal mutations

Outputs from clonal analysis can be found in **Table S10**.

#### Determining clonal VAFs and populations of duplex mutations from bulk WGS

Although duplex sequencing has an extremely low error rate, bWGS without the intrinsic error-correcting ability of duplex sequencing can still harbor several technical artifacts; thus, a pileup may contain an allele that resembles a duplex mutation at the same site but is actually a coincident sequencing or amplification error. Thus, we evaluated the pileup counts of BLT50 duplex mutations in bWGS from tissue homogenates’ bWGS, which did not come from the COLO829 donor and are thus highly unlikely to harbor most of the same somatic mutations as the BLT50 mixture. For each assay, we developed a pileup threshold that maximizes the number of BLT50 duplex mutations with pileups in COLO829 bWGS (the “pileup sensitivity”) and minimizes the number with pileups in non-COLO829 donors (the “pileup false positive rate”). At the maximum area under the receiver-operator characteristic curve (AUROC), the pileup sensitivity is ∼75% for Illumina-based duplex assays and 92% for Ultima ppm-seq assays. The false positive rates are comparable (5% for Ultima ppm-seq and ∼7% for Illumina-based) (**Figure S10A**).

For BLT50 duplex mutations that crossed our selected pileup threshold, the VAF distributions derived from bWGS of BLT50, tumor, and blood were highly multimodal and maintained a consistent structure across assays (**Figure S10B**). We hypothesized that each mode in the VAF distributions corresponds to a different clonal population captured within the BLT50 duplex mutations. Furthermore, given how the tumor and blood populations were mixed to yield the BLT50 sample, we speculated that each mode in BLT50 should correspond to some combination of modes from tumor and blood, resembling the clonal structure of a complex tissue.

Thus, we fitted a beta-binomial mixture model to extract the modes and examine their relationships to each other. All assays’ mutation counts were fitted together in the same model, but a separate model was fitted for each bWGS’s pileup results. For example, all nine assays were analyzed in the same model for pileups in BLT50, but a separate model was fitted for all nine assays together in BL. We inferred component parameters using 100 iterations of expectation maximization (although parameter estimates converged around 75 iterations). For the expectation step in each iteration, we bootstrapped 5000 mutations from each assay to ensure that assays were represented equally in estimating mode parameters. Likelihoods were computed using mode parameters for all mutations in each iteration’s maximization step. Thus, the estimated modes remain robust to the broad variation in mutation counts across assays.

## Supplementary Tables

**Table S1**. Summary of analysis metrics

**Table S2**. HDP-extracted signatures (excluding ppmSeq)

**Table S3**. MuSiCal-extracted signatures (excluding ppmSeq)

**Table S4**. Cosine similarities between HDP and MuSiCal signatures

**Table S5**. COSMIC decomposition of MuSiCal signatures

**Table S6**. MuSiCal signature exposures

**Table S7**. HDP signature exposures

**Table S8**. HDP-extracted signatures (including ppmSeq)

**Table S9**. MuSiCal-extracted signatures (including ppmSeq)

**Table S10**. Overlap of clonal mutations with ultra-deep bulk sequencing

## Supplementary Figures

**Supplementary Figure 1.**
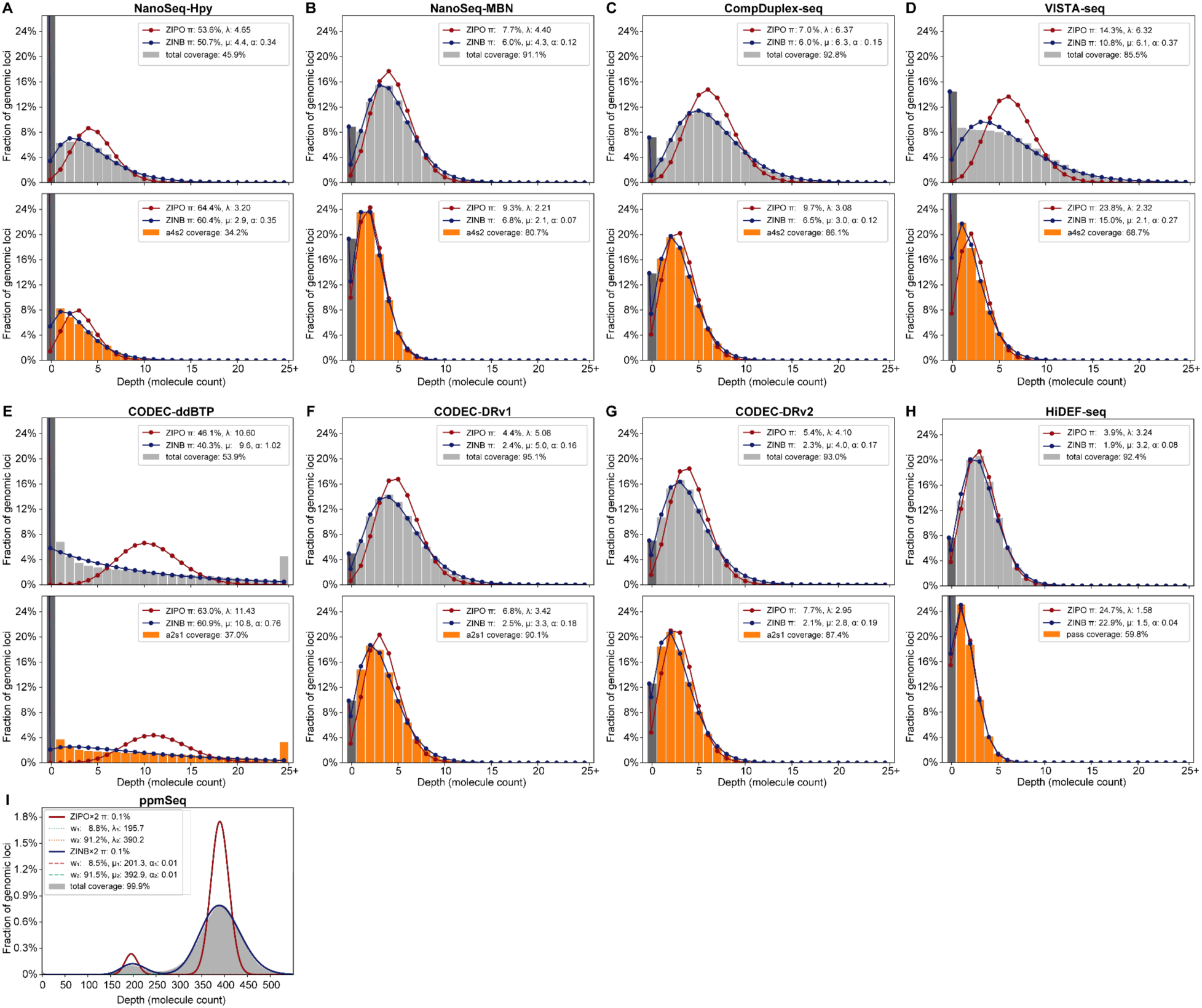
Modeling genome-wide duplex coverage distribution in COLO829 across NanoSeq technologies. For each library design, histograms show the empirical per-locus duplex coverage distribution for total molecules (gray) and interrogated base pairs (orange). Fitted curves represent Zero-Inflated Poisson (ZIPO; red) and Zero-Inflated Negative Binomial (ZINB; blue) models. Model parameters (π, μ, α) are displayed within each panel. π represents the proportion of structurally unsequenced loci, μ the mean duplex depth, and α the over-dispersion parameter (with α→0 corresponding to the Poisson limit). Across all technologies, ZINB consistently provides better fit to empirical data, capturing both the excess of zero-covered loci and the heavy-tailed depth distribution that arises from sequence-context and fragmentation biases.Total and high-fidelity coverages are indicated as percentages of the callable genome.

**Supplementary Figure 2.**
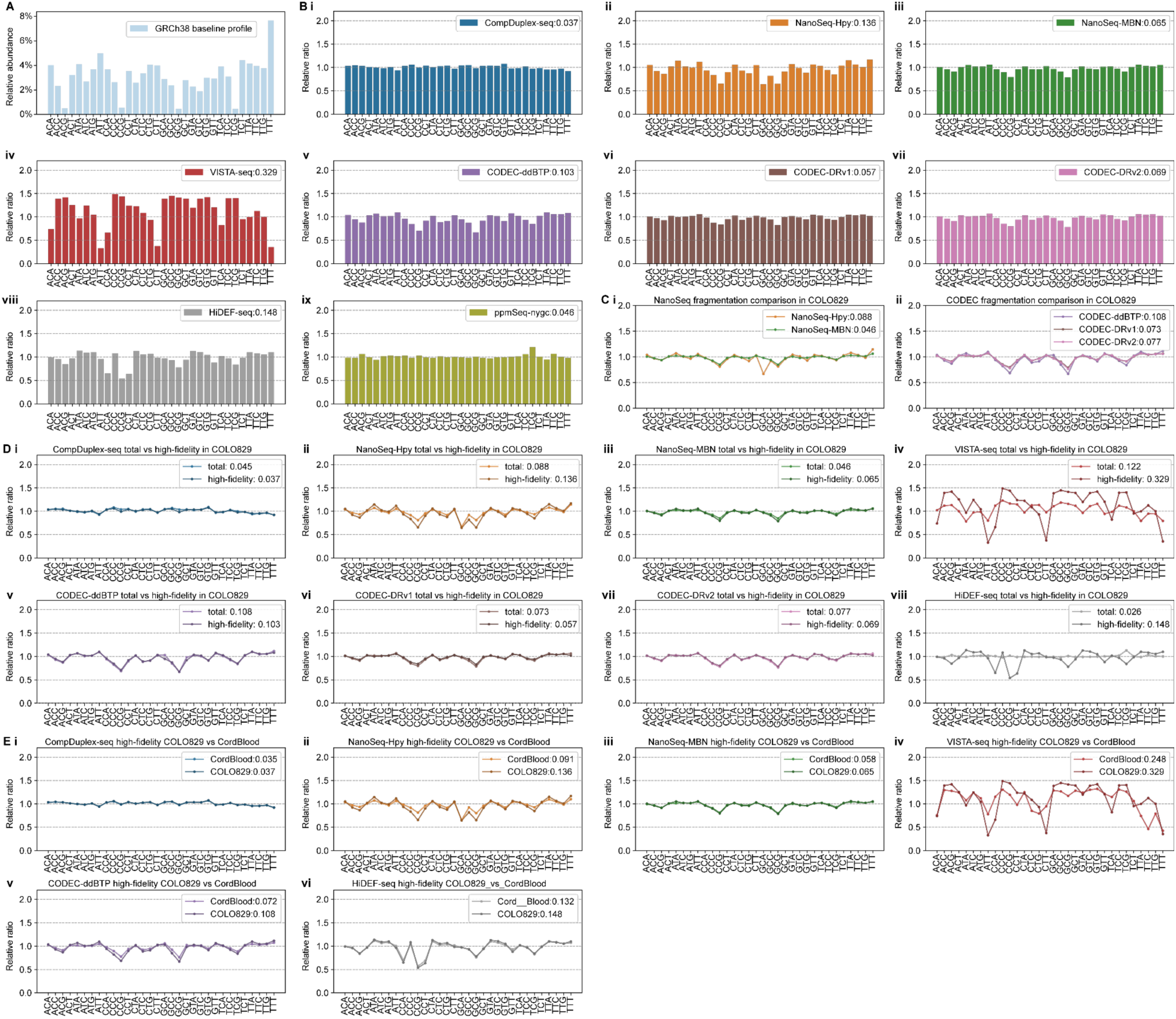
Trinucleotide Context Enrichment.X-axis: pyrimidine-centric trinucleotide contexts (32 classes). (A) Baseline trinucleotide composition of GRCh38 after excluding masked regions. (B) Per-library enrichment relative to (A): the ratio of each library’s trinucleotide abundance to the GRCh38 baseline. Numbers in the legend indicate the standard deviation across contexts and serve as a scalar measure of bias magnitude. (C) Effect of fragmentation strategy on trinucleotide bias, contrasting restriction-enzyme–based versus random fragmentation. (D) Effect of fidelity filtering on trinucleotide bias: “total” denotes unfiltered molecules, whereas “high-fidelity” denotes interrogated base pairs. (E) Effect of DNA material on trinucleotide bias, comparing Cord Blood with COLO829-BLT50 tumor spike-in.

**Supplementary Figure 3.**
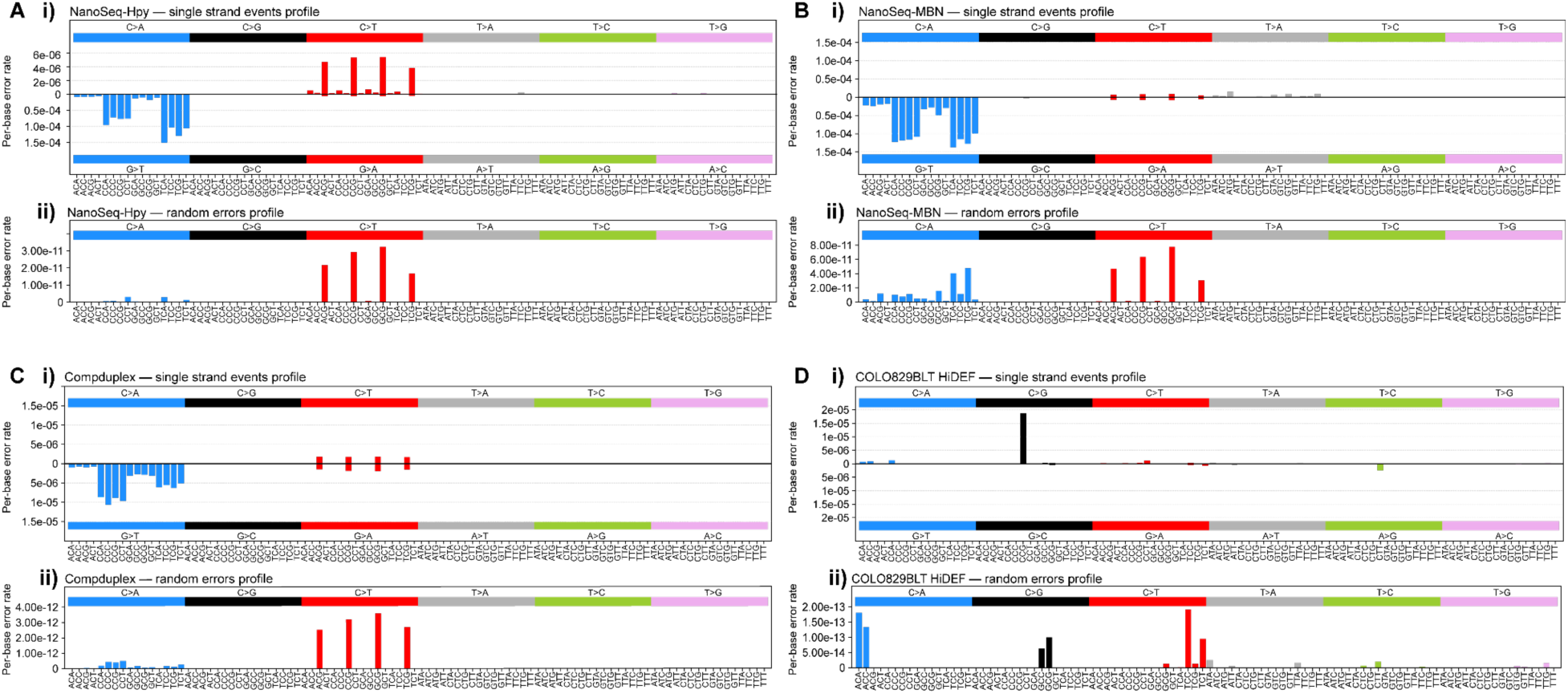
Single-strand events and random error rates. For each technology, (i) displays per-trinucleotide, per-strand potentials 𝑝_++_(𝑘, 𝑠). The x-axis follows the 96-class, pyrimidine-centric substitution order: above the axis are pyrimidine-centric single-strand events; below the axis are the complementary purine-centric single-strand events (reverse-complement pairs). Frequencies are normalized by the corresponding opportunities (interrogated base-pairs × 2; see Interrogated Base Pairs). Across methods, single-strand events are asymmetrically distributed across complementary contexts. (ii) shows the random error profile, computed as the independence product of complementary single-strand event potentials weighted by the library’s trinucleotide composition (see Trinucleotide Context Enrichment). This asymmetry in panel (i) leads to a reduced random-coincidence rate in panel (ii). Note that these values quantify the random-coincidence floor, not the full false-positive burden, which also includes systematic and context-coupled errors and depends on polymerase damage detectability.

**Supplementary Figure 4.**
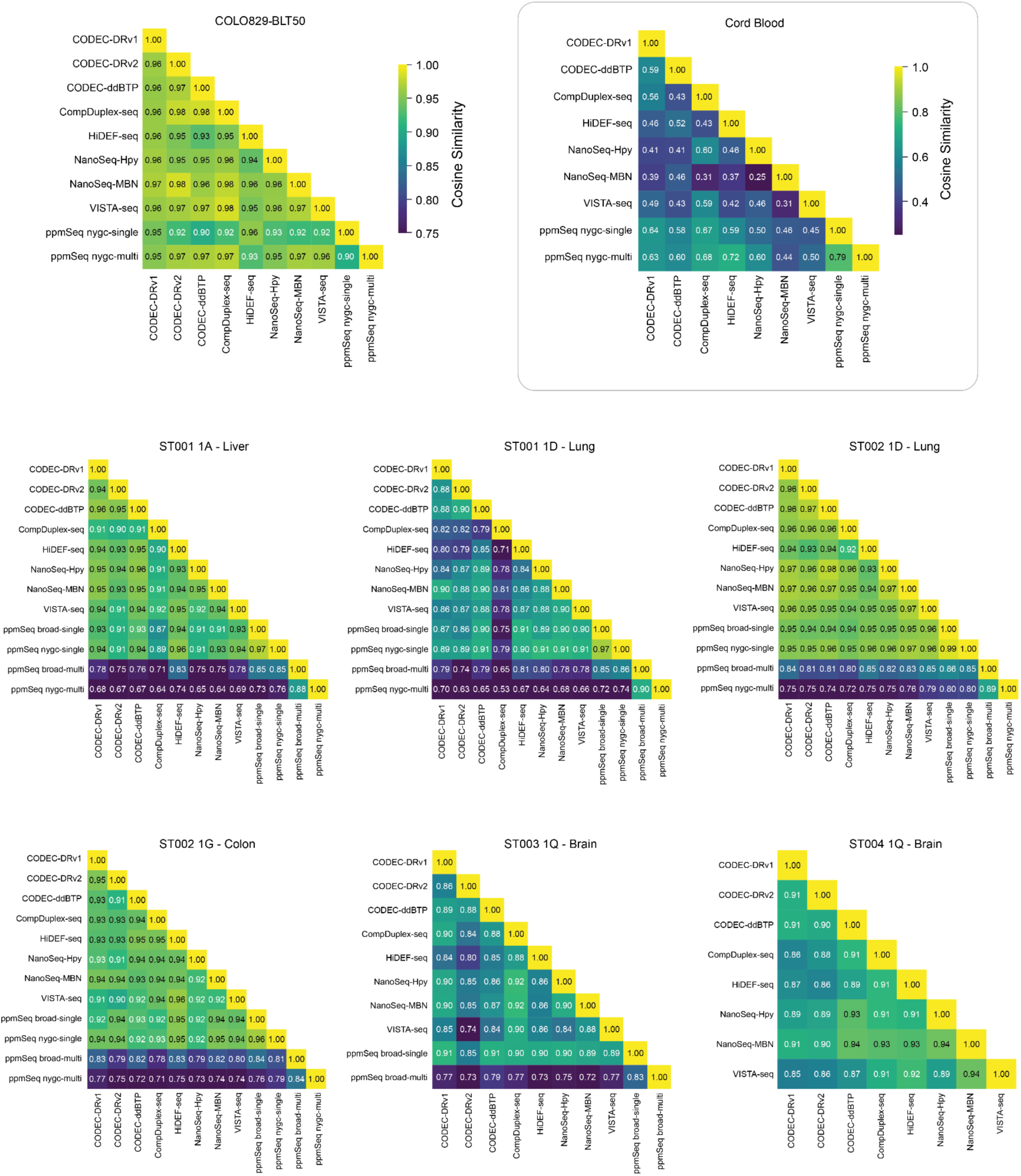
C**r**oss**-technology similarity of mutational profiles.** Cosine similarity heatmaps showing pairwise similarities between raw mutational spectra obtained from each sequencing technology, stratified by tissue and donor.

**Supplementary Figure 5.**
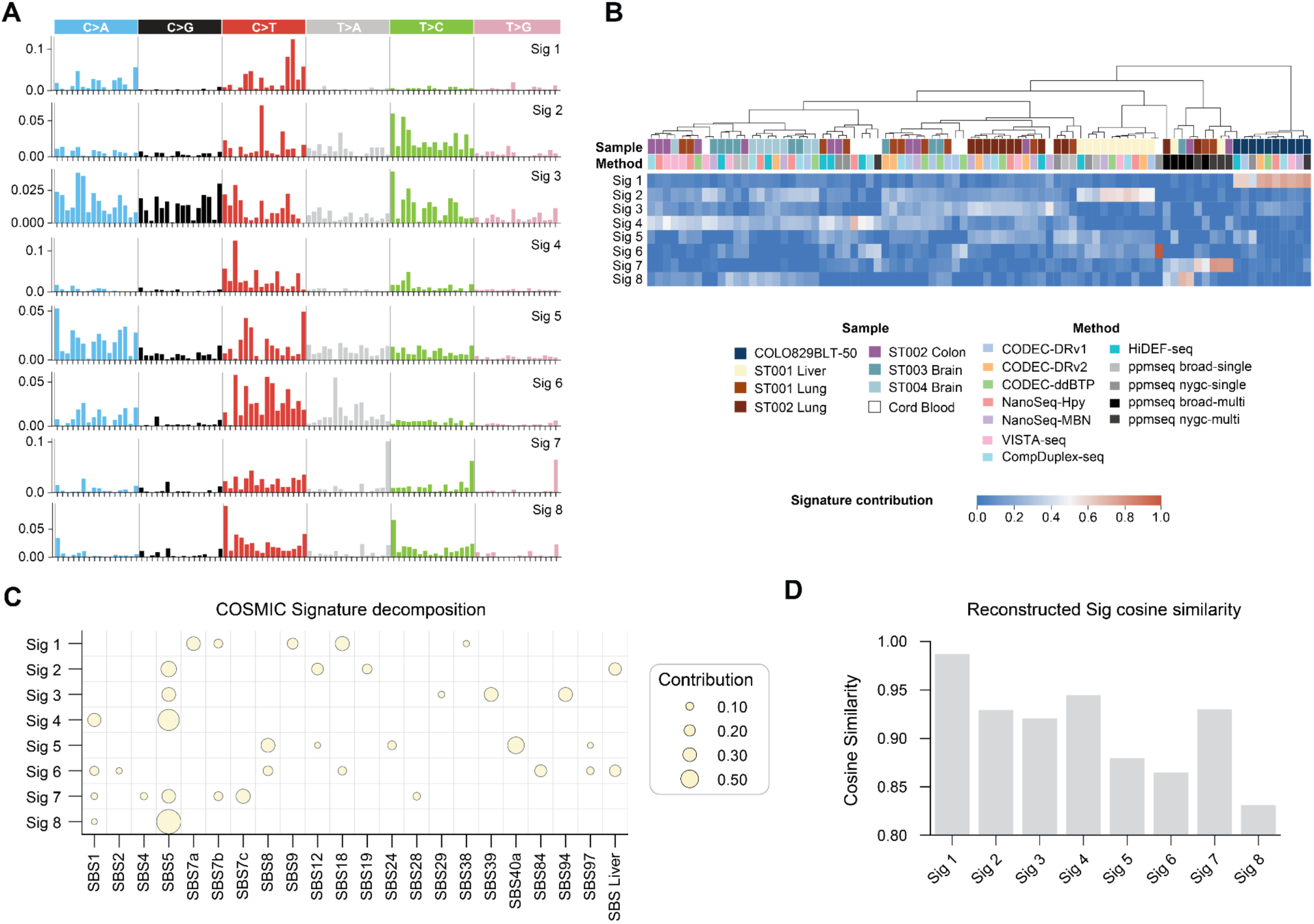
De novo mutational signatures extracted from the complete dataset, including all ppmSeq calls. **A**. Signatures extracted using NMF as implemented in MuSiCal. **B**. Heatmap of signature exposures across samples, hierarchically clustered by Ward’s method; top rows indicate sample, tissue, and sequencing platform. Signatures 6–8 appear ppmSeq-specific. **C**. Decomposition of de novo signatures into known COSMIC signatures; circle size indicates contribution proportion. **D**. Bar plot showing cosine similarity of each de novo signature to its reconstruction from top COSMIC signatures.

**Supplementary Figure 6.**
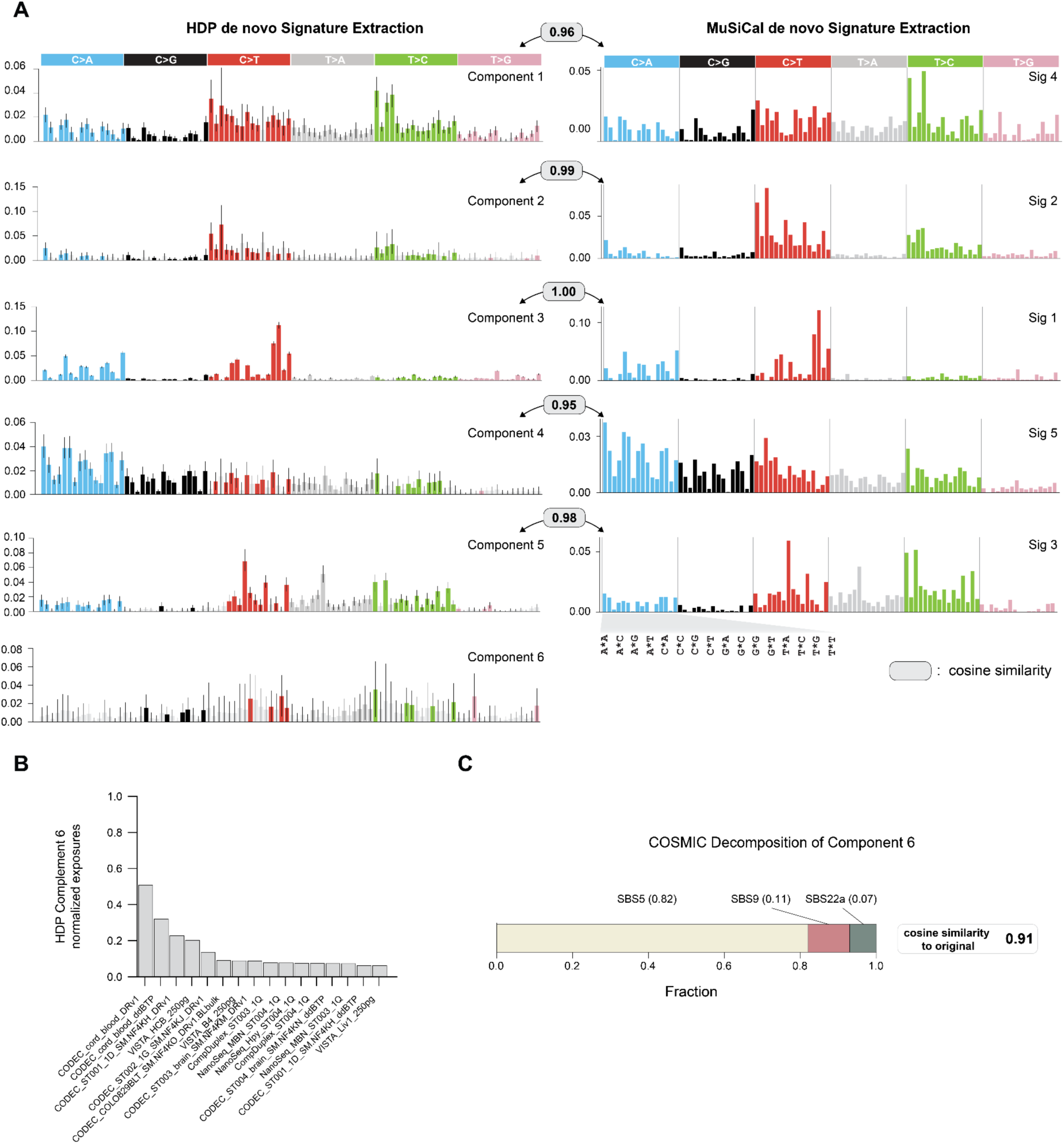
Comparative analysis of de novo mutational signatures identified by MuSiCal and HDP **A.** De novo mutational signatures recovered by MuSiCal and HDP. Grey boxes indicate cosine similarity between corresponding signatures. HDP identifies an additional component (Component 6). **B.** Top 15 normalized exposures after refitting Component 6. **C.** Reconstruction of HDP’s Component 6 from COSMIC reference signatures. Cosine similarity between the reconstructed and original de novo signature is shown.

**Supplementary Figure 7.**
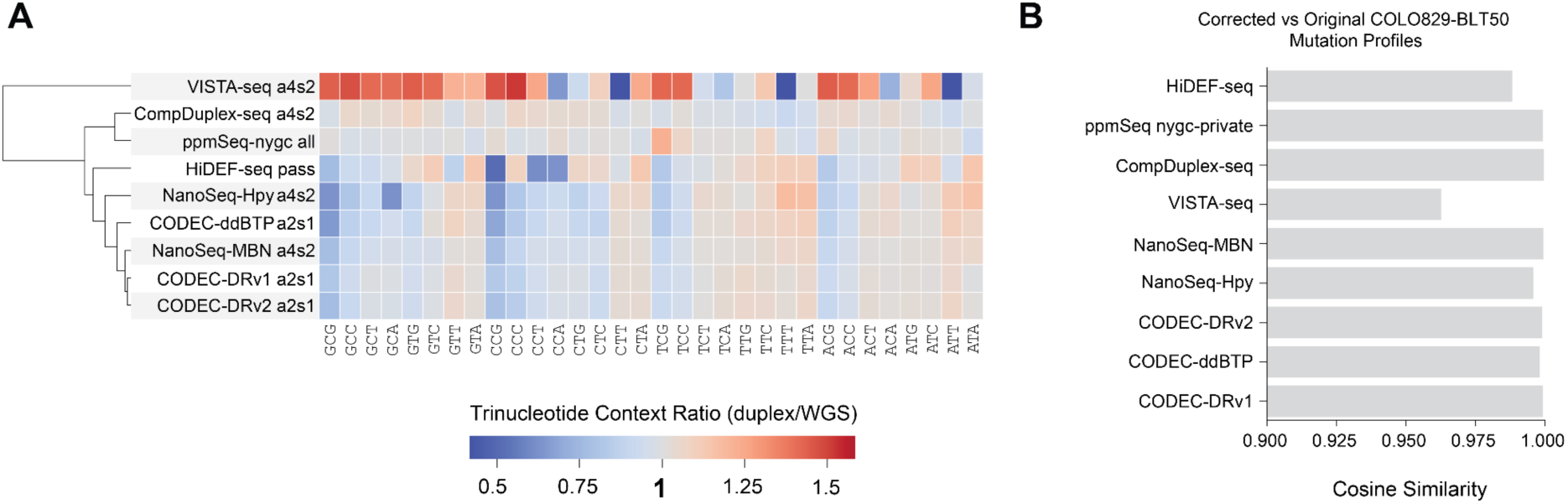
Influence of sequencing coverage bias in duplex technologies on COLO829-BLT50 mutational profiles **A.** Clustered heatmap showing the ratio of trinucleotide context abundances between each duplex sequencing assay and the reference genome (WGS, hg38). Assays are clustered to highlight similarities and differences in sequence bias across technologies. Per-technology trinucleotide biases inform correction of obtained mutational profiles (see Methods). **B.** Cosine similarity between corrected and original COLO829-BLT50 mutation profiles, illustrating the limited impact of tri-nucleotide sequence bias on the mutational profile as quantified by different duplex technologies for this sample.

**Supplementary Figure 8:**
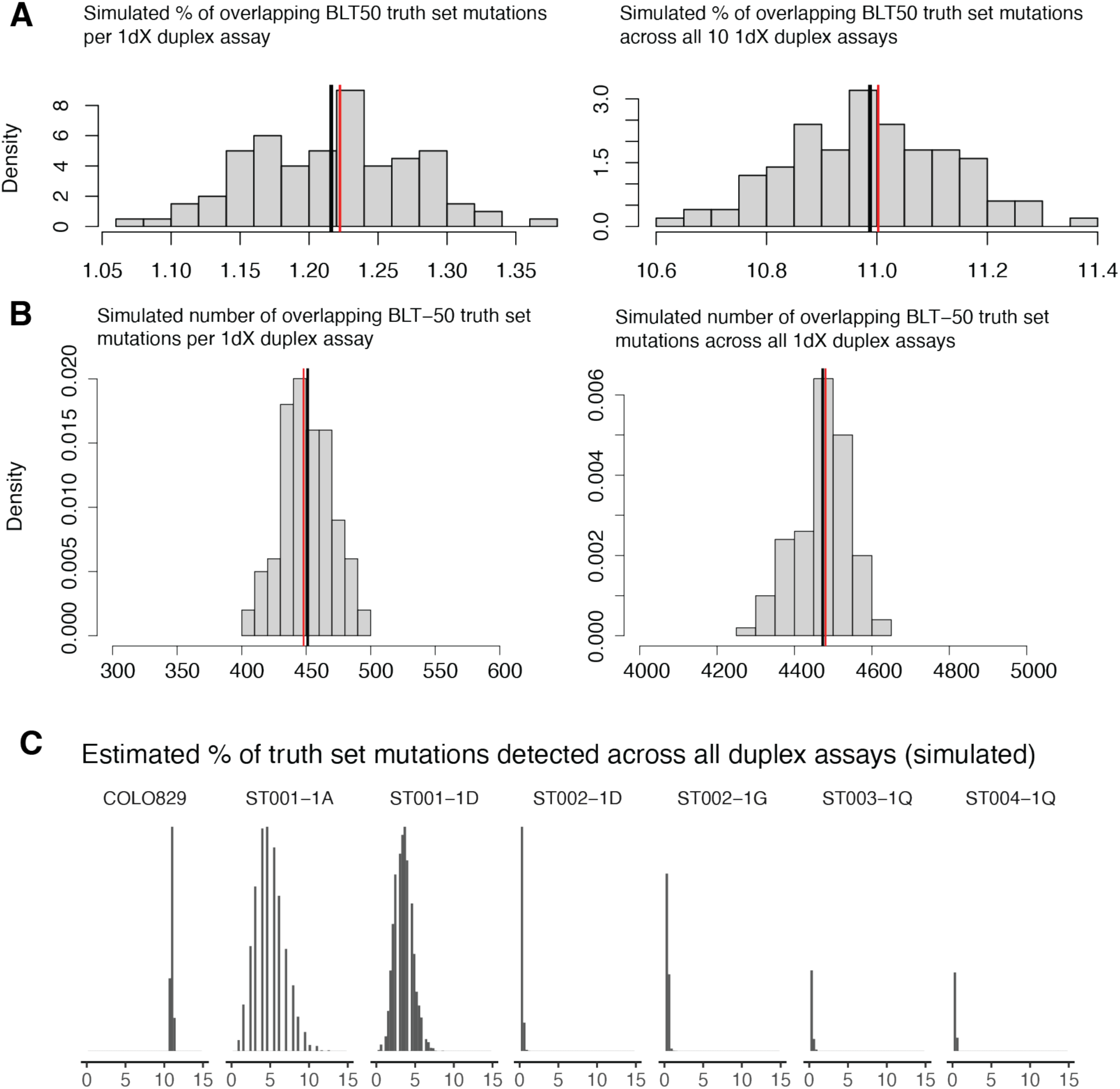
Simulated re-capture of clonal reference truth set mutations from COLO829-BLT50 and benchmark primary tissue samples. A). Simulated percent of BLT50 truth-set mutations that would overlap with mutations called from duplex sequencing at different units of duplex coverage. Left: 1 unit of duplex coverage (1dX). Right: 10 dX, or the aggregate of all shallow duplex sequencing datasets used in the benchmark. **B).** Simulated number of BLT50 truth-set mutations that would overlap with mutations called from duplex sequencing at different units of duplex coverage (left: 1 dX; right: 10dX). **C).** Estimated percentage of truth set mutations expected to be detected across all shallow duplex assays.

**Supplementary Figure 9:**
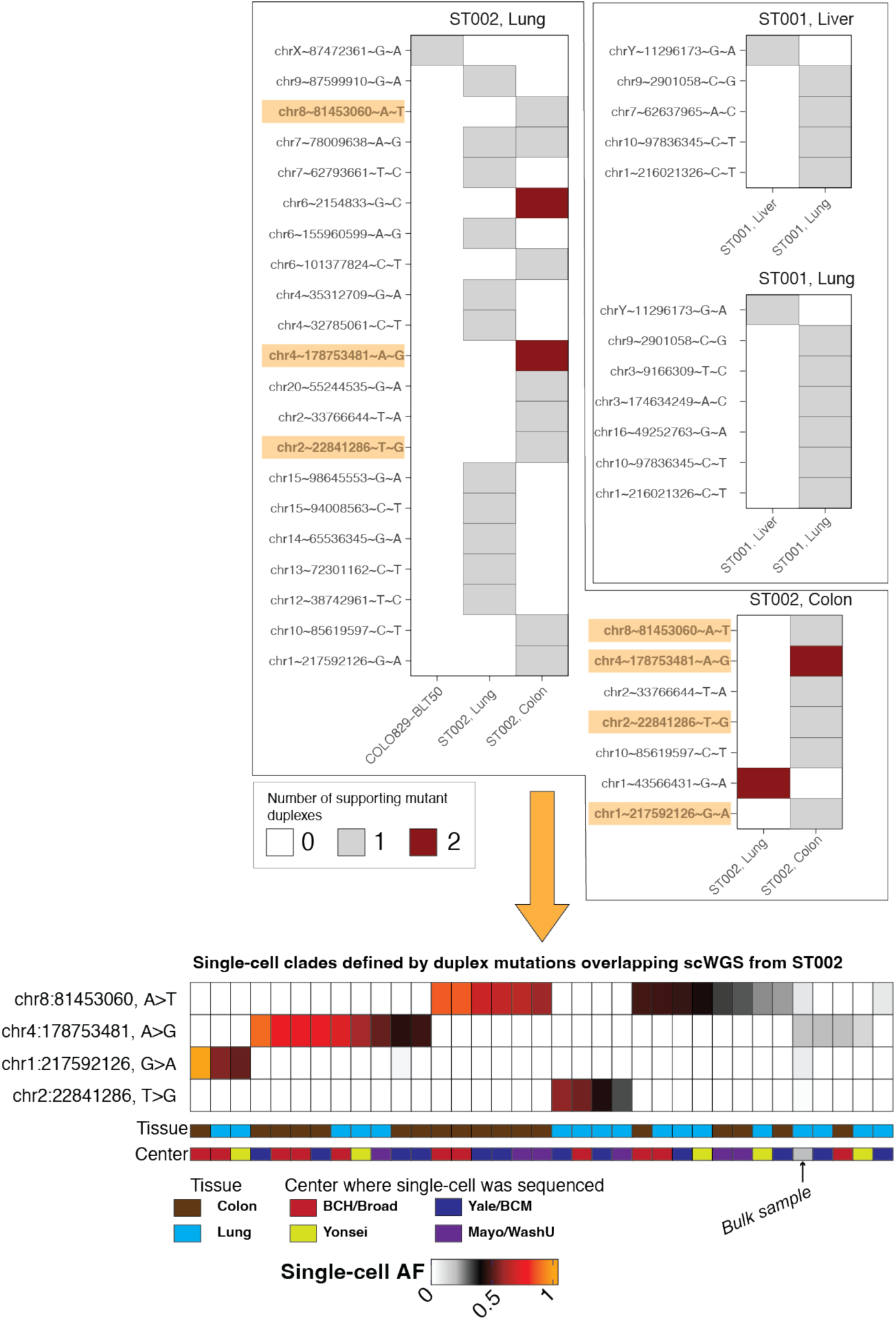
Overlap of duplex mutations across multiple primary tissue samples in the same benchmark donor. Top: Duplex mutations from benchmark primary tissues that overlapped any of the benchmark truth sets are shown. Heatmaps are organized by the donor where the duplex mutations originate; columns correspond to the truth set in which the mutation overlaps. Cells are filled by the number of supporting duplexes for the mutation. Bottom: A subset of duplex mutations from ST002 (highlighted) also overlapped mutation calls made from whole-genome sequencing of single cells ST002; cells are organized into clades defined by these duplex mutations.

**Supplementary Figure 10:**
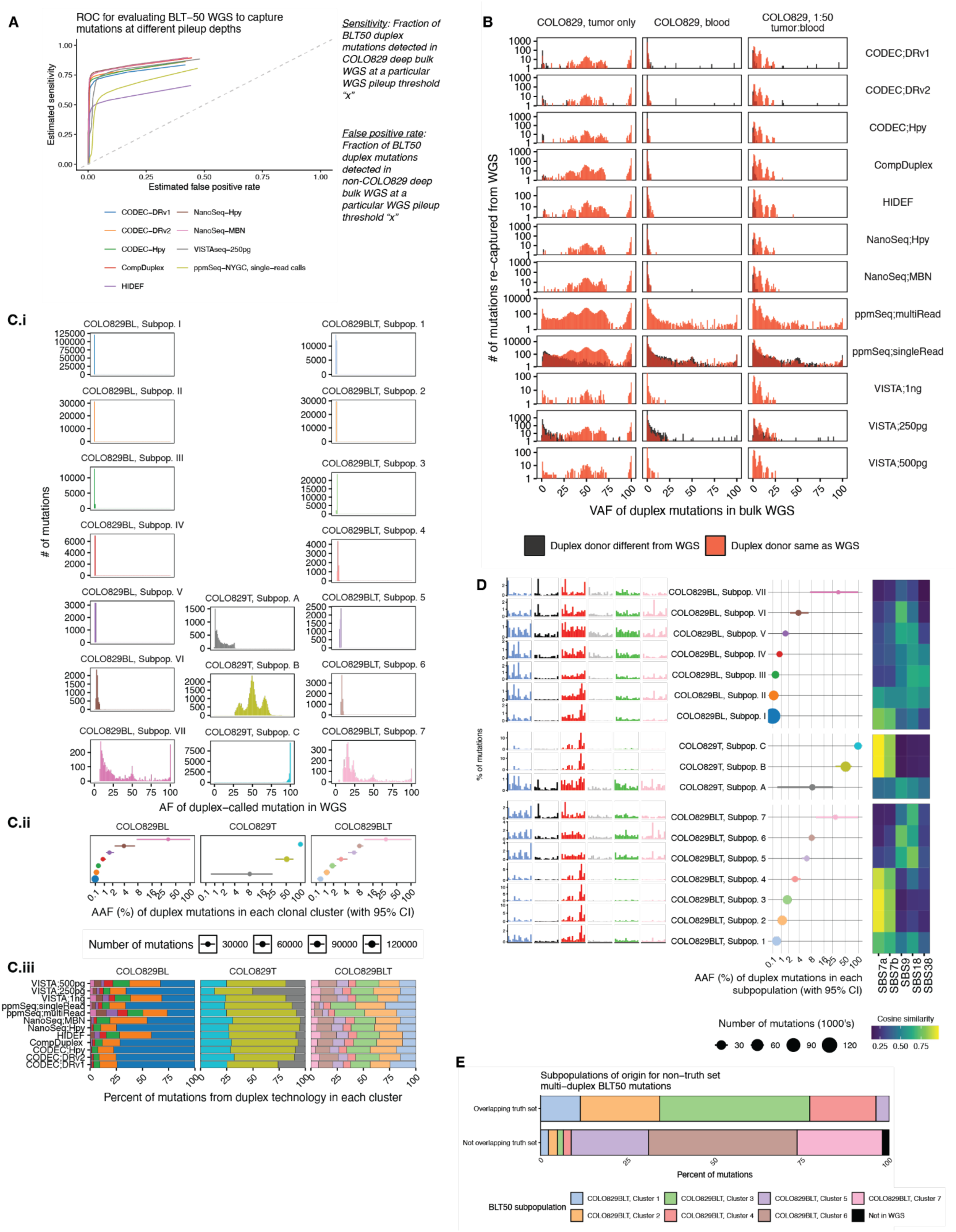
Detection and properties of clonal subpopulations in blood, tumor, and blood/tumor mixture as inferred from integrating duplex sequencing and deep **bulk WGS. A).** Receiver-operator characteristic (ROC) curves describing the sensitivity and specificity of using read pileups from bulk WGS (bWGS) of BLT50 (at different read level thresholds) to identify a mutation called from duplex sequencing of BLT50 as clonal. The sensitivity of a pileup is the fraction of duplex calls that are detected using bWGS pileups from the same donor. The false positive rate of a pileup is the fraction of duplex calls that are detected using bWGS from a different donor, suggesting that the bWGS pileup might comprise sequencing errors mimicking the duplex-called mutant allele. Different ROC’s were calculated based on duplex calls from different technologies. For each technology, the pileup threshold that maximizes sensitivity and minimizes the false positive rate was chosen to identify clonal mutations from duplex callsets. **B).** VAF spectra of duplex-called mutations based on pileups from the bWGS of BLT50 (right) and its individual tumor (left) and blood (middle) components. The VAFs of duplex mutations from BLT50 are shown in red; the VAFs of non-BLT50 duplex mutations but with alt alleles detected in the pileups of BLT50 and its respective components are shown in black. **C).** The different subpopulations identified by fitting a beta-binomial mixture model on the VAF distributions aggregated across duplex technologies (and normalized for duplex callset size) within each bWGS distribution. **Ci).** VAF distributions of individual subpopulations. **Cii).** Mean VAF and 95% confidence interval for each component. **Ciii).** Percentage of BLT50 duplex mutations from each technology assigned to each subpopulation. **D).** For each BLT50, tumor, or blood subpopulation are presented with its trinucleotide spectrum, mean VAFs (with 95% CI), and cosine similarities of spectra to COSMIC fitted components. **E).** The percentage of multi-duplex mutations (**Figure 5B**) that were assigned to different BLT50 clonal subpopulations is presented and separated by whether the mutation overlaps the BLT50 truth set.

